# Maize and wild relatives show distinct patterns of gene loss following polyploidy

**DOI:** 10.1101/2024.12.24.630189

**Authors:** Samantha J. Snodgrass, Margaret Woodhouse, Arun Seetharam, Michelle Stitzer, Beibei Liu, Matthew B. Hufford

**Author notes:** Corresponding authors: Samantha Snodgrass, Matthew Hufford.

## Abstract

Repeated cycles of polyploidy and gene loss during diploidization have shaped plant genomes over evolutionary time. This loss of gene sequence post-polyploidy is known as fractionation. Gene duplicates (*i.e.,* homoeologs) from each ancestral diploid genome (*i.e.*, subgenome) may fractionate differently, resulting in one subgenome being preferentially retained. Previous work suggests fractionation occurs shortly after a polyploidy event. Yet, few studies have explicitly compared genome-wide fractionation patterns across genera sharing the same whole genome duplication (WGD) to clarify how patterns of fractionation distinguish species over time. The Tripsacinae subtribe of grasses, which includes the genera *Tripsacum* and *Zea*, share an ancient allotetraploidy event ∼5-12 MYA. Using publicly available genome assemblies for the Tripsacinae subtribe including wild lineages and the domesticate maize, we investigate patterns of fractionation. Across genomes, the majority of fractionation is shared across species and one subgenome is preferentially retained. However, ∼35% of exons were fractionated differently across species, with significantly more differences between genomes of *Tripsacum* and *Zea*. We found no support of incomplete lineage sorting explaining fractionation differences across genera. Rather, the most striking trend is *Tripsacum* retaining a substantially greater proportion of homoeolog pairs than *Zea* where gene loss has been more prevalent.

## Introduction

Whole-genome duplication (WGD) instantly inflates gene content, duplicating every gene relative to diploid parental genomes, resulting in polyploidy. Despite repeated rounds of polyploidy (Wendel et al., 2016), the number of genes per genome is relatively similar across species (Choi et al., 2020). This indicates that, as polyploids rediploidize, gene loss or fractionation must be common. In the polyploid genome, a gene’s descendant, duplicate copies are referred to as homoeologs. While they begin as a pair after the WGD, one or both homoeologs may be fractionated over time. Homoeologs from one subgenome may be preferentially lost, a phenomenon known as biased fractionation (Thomas et al., 2006, Wendel et al., 2018, Li et al., 2021), which may be related to repeat content differences between genomes (McRae et al., 2022). However, the direction of that bias can change in some plant lineages (Ma et al., 2024).

Within the Andropogoneae tribe of grasses is the Tripsacinae subtribe that shares an ancient whole genome duplication that occurred approximately 5 – 12 million years ago (Gaut & Doebley, 1997; Schnable et al., 2011; Swigoňova et al., 2004; Woodhouse et al., 2010). The subtribe consists of two genera, *Tripsacum* (haploid chromosome n=18, Anderson 1944, Maguire 1961) and *Zea* (haploid chromosome n=10), which includes the crop, maize (*Zea mays* ssp. *mays*). Almost all *Zea* species and roughly half of *Tripsacum* species are cytologically diploid, with all current polyploids the product of subsequent rounds of WGD (Cutler and Anderson, 1941, Li et al., 1999, Pasupuleti and Galinat 1982). The Tripsacinae polyploidy event occurred through the hybridization of an extinct species similar to modern *Urelytrum* and *Vossia* and another similar to modern *Rottboellia-Hemarthria* (McKain et al., 2018). Alignment of any diploid *Zea* species to close unduplicated relatives, such as sorghum, results in two orthologous, syntenic regions from the Tripsacinae genome aligned to one region in the unduplicated species (Schnable et. al. 2011; Lovell et. al. 2022). *Zea* species show strong synteny and collinearity to each other but extensive chromosomal rearrangements when aligned to other grasses (Schnable et. al. 2011; Chia et. al. 2012; Lovell et. al. 2022). The extent of synteny between *Tripsacum* and *Zea* has not been analyzed via comparative genomics (but see Galinat 1973). Therefore, while previous studies predict that fractionation patterns will be similar (*i.e.,* which homoeologs are lost and amount of loss) between descendents of the same WGD event, this has yet to be measured directly in the Tripsacinae (Gault et al., 2018; Sankoff et al., 2010).

Fractionation studies of this subtribe have focused on the domesticate, maize, finding most homoeologs have either been reduced to single copy or experienced partial loss across genomes (∼43% and ∼16% respectively, Hufford et al., 2021). Biased fractionation by small deletion events identified in *Zea mays* ssp. *mays* B73 (Zm B73) affected maize subgenome 2 (M2) more than maize subgenome 1 (M1) (Schnable et al., 2011; Woodhouse et al., 2010). SNPs in M1 regions of the maize genome show greater association with phenotypic variability across traits than M2 (Renny-Byfield et al., 2017) suggesting the M1 subgenome retains more function. Comparisons between two temperate maize genomes found little differential fractionation (*i.e.*, homoeologs of a gene are retained or lost in different ways in one genome versus another), suggesting fractionation had occurred before divergence of these lines (Brohammer et al., 2018). When expanded to include more diverse maize assemblies, differences in retention and loss were greater between more diverged lines (Hufford et al., 2021). Fractionation specific within subsets of lineages contrasts with the prevailing hypothesis that all fractionation occurs shortly after a WGD event and should thus be the same across all descendents (Schnable et al., 2011). Rapid loss of homoeologs has been observed in the aftermath of other WGD events across a wide breadth of systems, including angiosperms (Li et al., 2016), *Brassica napus* (Szadkowski et al., 2010), *Paramecium*, *Arabidopsis*, vertebrates, yeast, and teleosts (Sankoff et al., 2010). Yet in other lineages like *Brachypodium*, homoeologs have been retained for long periods of time and show gradual loss (Scarlett et al., 2023). Given limited genomic resources for the wild species of the subtribe, patterns of fractionation and their relationship to lineage divergence within the Tripsacinae have not been studied.

This study uses the recently published pseudomolecule assemblies of wild *Zea* species and *Tripsacum* along with previously published diverse maize nested association mapping (NAM) genomes to investigate the patterns of exon fractionation relative to diploid relatives, *Sorghum bicolor* and *Anatherum virginicum* (Andropogoninae, divergence ∼10MYA), in the Tripsacinae (diverged from Andropogoninae ∼19MYA, Stitzer et al., 2025; Hufford et al 2021). By including the wild Tripsacinae species, we investigate fractionation at larger divergence times than between any two maize accessions (∼650 KYA between *Tripsacum* and *Zea* to ∼67 KYA between *Zea mays* subspecies, Chen et al 2022). We identified fractionation relative to a conserved set of gene annotations from sorghum through whole genome alignments and characterize the deletions underlying sequence loss. We examined patterns of retention or loss of exons and genes between subgenomes as well as the status of homoeologous pairs across genomes. This allowed us to identify instances where fractionation differed. Further, whole genome alignments to sorghum highlighted extreme differences in the chromosomal structural history of the genera. Based on these observations, we hypothesize a potential relationship between the process of fractionation and the rearrangement histories of ancestral chromosomes.

## Results

### Multiple, small deletions underlie most fractionation

We used 69,269 exons conserved between sorghum and *A. virginicum* as a diploid reference set to model genes and exons from the extinct diploid progenitor species of the Tripsacinae. While these species are not independent outgroups, they are the closest relatives with reference quality assemblies of the extinct diploid progenitor species of the Tripsacinae, and we anticipate exon gain to be rare at these divergence scales (Xu et al., 2012).

Deletions were commonly observed in coding sequences across both subgenomes and across all species. Most of these deletions were small (1-10 bp in length), though longer and overall more deletions were observed in the M2 subgenome compared to the M1 subgenome (Figure 1A). This matches higher overall mapping rates for M1 sequences (mean ∼77%) versus M2 sequences (mean ∼64%). AnchorWave relies on genic, syntenic anchors to build alignments, which won’t occur in highly deleted and rearranged segments of the genome, resulting in less overall mapping for the more fractionated subgenome (Supp. Figure 2). Extensive deletions correspond to an M1 homoeolog missing between 20-23% and an M2 homoeolog missing 30-35% of its coding sequence on average. Further, both *Tripsacum* genomes show greater coding sequence retention across both subgenomes compared to *Zea* genomes (Figure 1B). For a given exon, multiple segregating and overlapping deletions can occur. The percentage of deletions nested in other deletions is high across both subgenomes (∼30-50%) while the percentage of deletions where an exon intersects with only a single deletion is < 10%. More than 67% of exonic deletions were smaller than the exon itself, but around 25% of deletions intersected with multiple exons due to their large size. Of those deletions that occur within a single exon, we asked how many of those deletions result in a frameshift, here meaning a deletion length that is not a multiple of three. We observed that exons with frameshift deletions versus those without frameshift deletions occurred at similar levels in homoeologs from subgenome M1, but there were more frameshifted exons in homoeologs from subgenome M2 (Figure 1C). From this point, we define an exon as fractionated if any deletion overlaps the exon’s boundaries.

**Figure 1.**
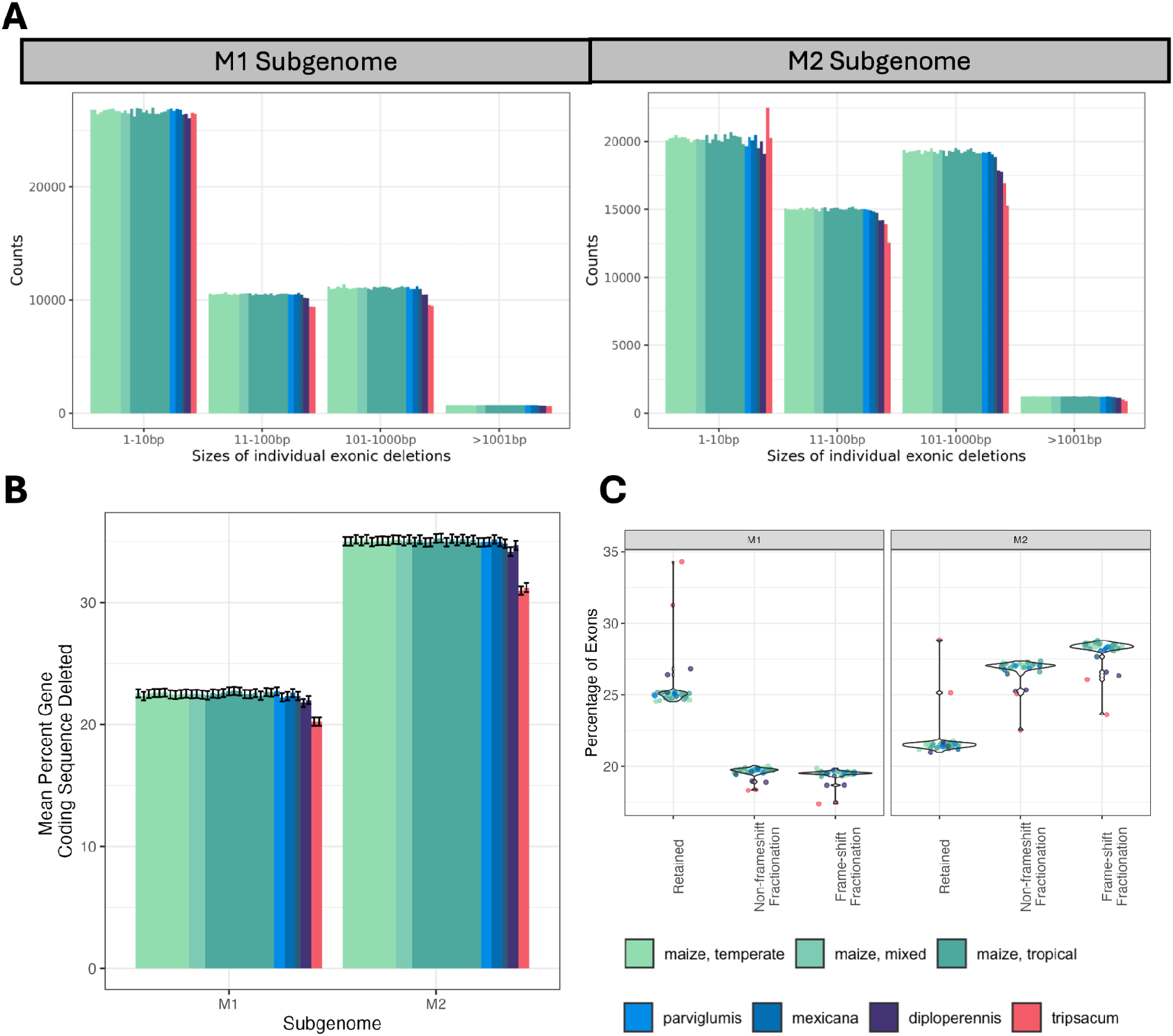
Deletion Characterization. (A) Counts of exonic deletions per genome by subgenome. Each bar represents one genome colored by broad taxonomic groups according to the key at bottom of figure (same key used across all panels). Bar height represents the count of deletions relative to sorghum in each size class by basepairs. (B) Average coding sequence lost per gene by subgenome. Each bar represents one genome colored by broad taxonomic groups. Bar height represents the mean average coding sequence lost per gene with error bars representing 1 standard error. (C) Percent of total exons that are either retained, fractionated without a frameshifting deletion, or fractionated with a frameshifting deletion. Points represent individual genomes colored by broad taxonomic groups and violins show overall distribution across all genomes.

**Figure 2.**
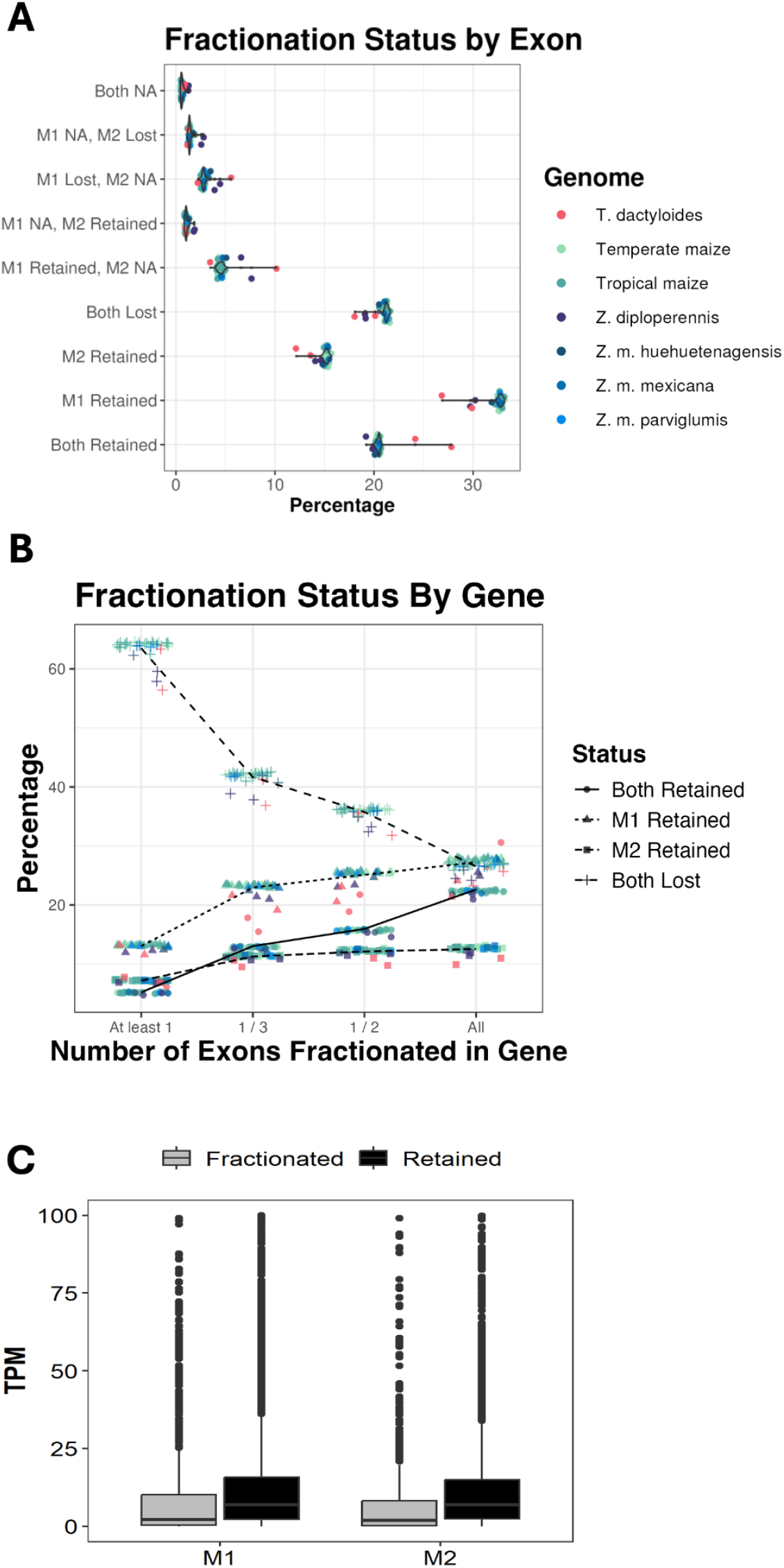
Fractionation status of homoeologous pairs. (A) Fractionation status of exon pairs. Points represent individual genomes and violins represent overall distribution across genomes. Statuses with “NA” represent an exon that was not alignable and treated as missing data. Color is by broad taxonomic groups. (B) Fractionation status of gene pairs across various thresholds. Points represent individual genomes. Shape corresponds to status (circle: both retained, triangle: M1 retained, square: M2 retained, cross: both lost). Color is by broad taxonomic groups. Lines intersect the mean for each status at each threshold. (C) Expression level of genes in maize leaves based on fractionation status across each subgenome.

### *Tripsacum* retains more exons and genes than *Zea*

Of the 69,269 reference exons to which sequences could be aligned, over 50% are retained as single copies. The remaining exons are evenly split between both copies being either retained or lost (Figure 2A, Table 1, 2, Supp. Table 1, 2). Unalignable exons cause a missing fractionation status in less than 10% of calls on average across genomes. *Tripsacum dactyloides* genomes (Td FL and Td KS) have more retained exons and better alignment to sorghum than do the *Zea* genomes (mean mapping percent *Tripsacum* M1 = 80%, M2 = 66%, *Zea* M1 = 77%, M2 =63%, Supp. Figure 2). Across all genomes, the M1 homoeologous exon is more often retained than the M2 homoeologous exon (Figure 2A, Supp. Figure 3). This supports the documented biased fractionation of M2 (Schnable et al 2011, Hufford et al 2021).

**Figure 3.**
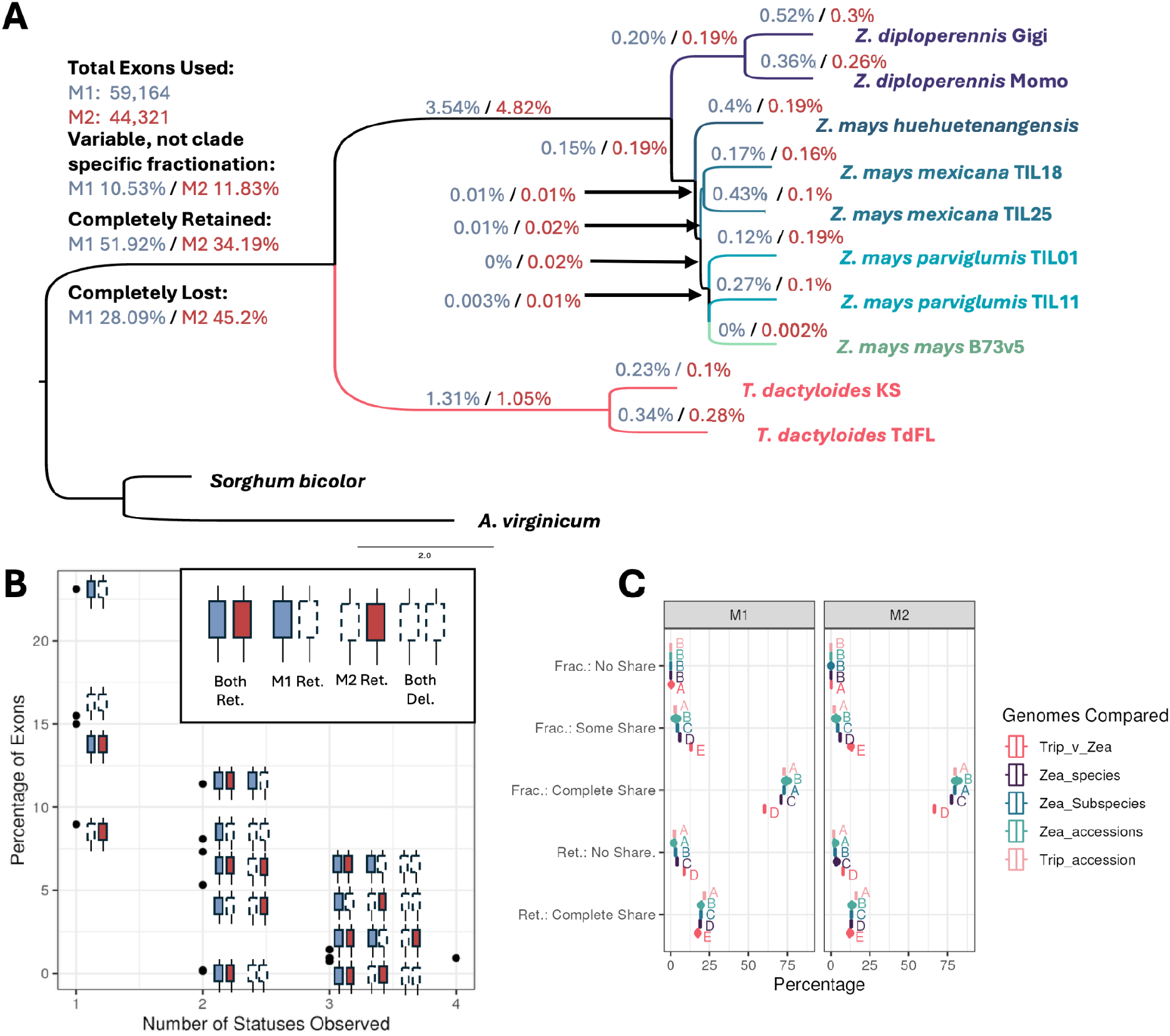
Sharing of fractionation across genomes. (A) Sharing of exon fractionation along the phylogeny from Stitzer et al. (2024). Exons must be aligned across all genomes to be used. Each branch is labeled with percent of exons in each subgenome that have been fractionated in every descendent lineage of the branch and retained in all other lineages (monophyletic fractionation). Scale bar is substitutions per site. (B) The number of statuses observed for a single homoeologous pair of exons across all genomes. Exon pairs are not required to be aligned across all genomes to be included. Black points indicate the percentage of all exons observed per category and the set of statuses observed in that point are illustrated to the right of each point, except for 4 which represents all statuses were observed. (C) Distributions of pairwise comparisons of exon loss sharing between two genomes. Genes that show either complete sharing or no sharing due to gene retention (categories starting with “Ret.”) and genes showing either complete, some, or no sharing of fractionation (categories starting with “Frac.”). Boxplots are colored by the broad relatedness of the genomes being compared. Letters indicate significant differences between comparison levels for each of the five categories by Tukey post-hoc tests.

**Table 1.** Counts of reference exons of each fractionation status across genomes shown in Fig 4. The total number of reference exons is 69,269.

| **Genome** | **Both Retained** | **M1 Retained** | **M2 Retained** | **Both Lost** | **M1 Retained, M2 NA** | **M1 NA, M2 Retained** | **M1 Lost, M2 NA** | **M1 NA, M2 Lost** | **Both NA** |
| --- | --- | --- | --- | --- | --- | --- | --- | --- | --- |
| Zm B73 | 14104 | 22862 | 10614 | 14807 | 2994 | 717 | 1897 | 859 | 415 |
| Zv TIL11 | 14068 | 22672 | 10542 | 14743 | 3206 | 749 | 1963 | 824 | 502 |
| Zd Gigi | 13042 | 20019 | 9648 | 12593 | 5528 | 1716 | 3473 | 2245 | 1005 |
| Td FL | 19267 | 20703 | 9415 | 13932 | 2387 | 701 | 1523 | 797 | 544 |
| Td KS | 16735 | 18596 | 8392 | 12497 | 7027 | 679 | 3853 | 878 | 612 |

**Table 2.** Average counts of reference exons of each fractionation status across all genomes by broad taxonomic grouping. The total number of reference exons is 69,269

| **Taxon** | **Both Retained** | **M1 Retained** | **M2 Retained** | **Both Lost** | **M1 Retained, M2 NA** | **M1 NA, M2 Retained** | **M1 Lost, M2 NA** | **M1 NA, M2 Lost** | **Both NA** |
| --- | --- | --- | --- | --- | --- | --- | --- | --- | --- |
| Maize (Zm) | 14174 | 22715 | 10535 | 14738 | 3107 | 709 | 1931 | 962 | 394 |
| Parviglumis (Zv) | 14144 | 22636 | 10457 | 14735 | 3212 | 746 | 1963 | 907 | 467 |
| Mexicana (Zx) | 14099 | 22619 | 10676 | 14563 | 3213 | 705 | 2054 | 923 | 415 |
| Huehuetenangensis (Zh) | 13916 | 22084 | 10267 | 14207 | 3519 | 906 | 2414 | 1202 | 754 |
| Diploperennis (Zd) | 13518 | 20761 | 9938 | 13243 | 4918 | 1277 | 2890 | 1850 | 871 |
| Tripsacum (Td) | 18001 | 19649 | 8903 | 13214 | 4707 | 690 | 2688 | 837 | 578 |

Gene fractionation was called across four different thresholds based on the proportion of exons fractionated within the gene model. These ranged from the least stringent (*i.e.,* at least one exon is fractionated) to the most stringent (*i.e.*, all exons are fractionated). We did this to see if the high percentage of homoeologs called fractionated was due solely to lenient cutoffs or fractionation of most or all exons. As expected, the number of both homoeologs called fractionated decreases as the cutoff stringency increases (Figure 2B). However even at the most stringent threshold, around 25% of homoeologs have been completely lost from both subgenomes and only around 12% of homoeologs are completely retained across both subgenomes. Thus, most fractionated homoeologs have fractionated most or all of their exons (Figure 2B). Across thresholds, single copy retained homoeologs were more often from the M1 subgenome than the M2 subgenome. Genic patterns of fractionation also mirrored the exon patterns where *Tripsacum* genomes retain more homoeologs as pairs than *Zea* genomes across all thresholds (Figure 2A, B).

Fractionation of homoeologs appears to correlate with transcriptional differences as well. Using the publicly available maize Nested Association Mapping (NAM) leaf RNAseq data (Hufford et al., 2021), we observed that across both subgenomes, retained homoeologs were more highly expressed than fractionated homeologs (Figure 2C). That fractionation correlates with a loss of gene expression suggests that fractionated genes may have lost some of their original function.

### Most fractionation is shared across all lineages or within genera

Considering exons that can be aligned across all genomes, we investigated how much fractionation was shared by phylogenetic relatedness. With almost 600KY since the divergence of *Tripsacum* and *Zea* (Chen et al., 2022), the majority of exons (∼80%) are either completely retained or fractionated for both M1 and M2 across all genomes (Figure 3A). However, M1 shows more complete retention than M2 (51.9% vs 34.2% respectively). Between genera, more exons are fractionated in *Zea* compared to *Tripsacum* genomes (3.54% vs. 1.31% in M1, 4.8% vs. 1.05% in M2). Moving towards the tips of the phylogeny, we find fewer instances where fractionation is either specific to a given genome (0.002-0.52%) or shared by a small monophyletic subset of genomes below the level of genus (1.31%-3.54%)(Figure 3A, Supp Figure 5).

About 10% of all aligned exons share fractionation across a group of non-monophyletic genomes or are discordant. Generally these are observed as either two non-sister lineages sharing fractionation of an exon or all but one lineage sharing fractionation of an exon (Figure 3A, Supp. Fig. 5). To see if this 10% might be due to incomplete lineage sorting (ILS), we estimated the probability of expected ILS for these lineages and compared these to observed instances of discordance. Our estimates of expected ILS range from close to zero to ∼65% depending upon parameters Ne and divergence time (Supp. Table 3, Supp. Figure 6). The greatest chances of expected ILS are between the subspecies of *Zea mays* (*mexicana, parviglumis,* and *mays*) given their recent divergence. Across all scenarios we investigated, observed discordance was much less than expected, likely because deletion variants are not neutral or nearly neutral. However, observed discordance between the most diverged lineages (scenario MVT, maize-parviglumis-tripsacum) was more common than the least diverged lineages (scenario MVX, maize-parviglumis-mexicana) for both subgenomes, opposite theoretical expectations (Supp. Fig 6C). Thus, while ILS may contribute to non-monophyly of fractionated exons, multiple mutational origins resulting in exon loss, especially between *Tripsacum* and *Zea,* likely underlie these observed discordance patterns.

Previous work has demonstrated that certain types of genes are more likely to be fractionated due to dosage balance of gene products (Birchler & Veitia, 2010) and create temporal patterns of loss and retention of certain gene classes (Barker et al., 2008). From these works, we expect that gene categories associated with essential functions are more likely to be shared in their retention or fractionation across all genomes while gene categories more associated with environmental responses would be more variable across genomes. To that end, we evaluated GO term enrichment for our subset of genes which share fractionation or retention across monophyletic groups of genomes.

For brevity, we summarize the GO terms with high fold enrichment (>2) or depletion (<0.5) though we report all significant GO terms (Supplemental Data 1). For genes which shared fractionation or retention across all genomes for either M1 or M2 subgenomes, often the same GO terms were enriched for retention and depleted for fractionation. These are often related to protein localization (to the vacuole for M1 and to membranes for M2), ribosomal subunits and functions, mitochondrial activity, and structural molecular activity, particularly actin for M1. M1 also showed conservation of genes in voltage gate channels, exosomes, and late endosomes. M2 showed additional conservation of ER vesicle transport and sulphur transferase activity. All together, these categories correspond to essential functions of living cells, as expected.

When considering genes that show fractionation or retention only across *Tripsacum* genomes or *Zea* genomes, both sets of M2 genes show enrichment of fractionated genes in categories associated with regulation of organelle organization. *Zea* genomes further showed enrichment of fractionated M2 genes associated with nuclear division. *Zea* genomes also show M1 fractionated genes enriched for RNA polymerase II and chromosome GO terms. These patterns suggest that fractionation or retention of certain gene classes may have differed between *Tripsacum* and *Zea*.

### Differential fractionation across genomes

While most fractionation is shared across genomes, we investigated instances in which retention or loss of an exon differed across genomes. We categorized differences by the number of fractionation statuses observed for a given pair of exons. For example, if a pair of exons in one genome are both retained while in another genome only one exon is retained, then those are two fractionation statuses observed. In line with previous work (Brohammer et al, 2018), we observe the majority of homoeologous exon pairs fall into a single status across all genomes (∼60%) (Figure 3B). As the number of fractionation statuses increases, there’s a precipitous drop in the percentage of exons observed. Genes which had more conserved exon fractionation status, *i.e.*, only a single status across genomes, were statistically shorter than genes with multiple fractionation status (∼500 bps or 1 exon shorter, p<2e-16 for both length and total exon count), but otherwise there were no biases in gene length across categories. Genes that are only observed with a single status across genomes showed significant GO term enrichment across a variety of biological, molecular, and cellular processes. Briefly, genes with exons “both lost” were enriched for transcription factor related terms; genes with exons “both retained” were enriched for terms related to protein binding, regulation, and modifications; genes with exons “M1 retained” were enriched for terms related to nuclease activity but depleted for terms related to transcription factor binding; and genes with exons “M2 retained” were enriched for catalytic activity and vesicle related terms ([Supplemental Data 1](https://docs.google.com/spreadsheets/d/1UxduvHWCERoCruozuRw2aa9VXgdSa4VTJoaBNIdefww/edit?usp=sharing)). Of those exon pairs observed with two fractionation statuses across genomes, ∼33% (7093 exon pairs) are differences between the *Tripsacum* accessions and *Zea* accessions. Exons with three and all observed statuses are a limited percentage of the total, ∼3% and ∼ 1% respectively (2,539 and 555 exons respectively). Genes which had exons within these differentially fractionated groups were not statistically enriched or depleted for GO terms, with one exception. Genes with exons observed with three statuses across genomes were enriched for cellular GO terms related to organellar and cytoplasmic membranes.

At the gene level, many homoeologs show loss of the same exons across all genomes (Supp. Figure 7, Supp. Figure 8). Yet, some homoeologs show fractionation of different exons across genomes. In pairwise comparisons of genomes, a homoeolog could have complete sharing or difference of fractionation status due to both or one being completely retained respectively. If fractionation occurred for exons of the homoeolog in both genomes, the fractionated exons could be all the same (“Completely Share”), some but not all the same (“Some Share”), or completely different (“No Share”). Homoeologs which only share some or none of the fractionated exons across genomes may provide evidence of multiple origins of fractionation and possibly convergence for a homoeolog towards fractionation. The vast majority of homoeologs (∼91%) across both M1 and M2 show complete sharing of fractionation across all genome pairs (Supp. Figure 8B,C). Significantly more homoeologs fractionate only some or completely different exons in pairwise comparisons between *Tripsacum* and *Zea* genomes (Figure 3C, Supp. Figure 8A,D,E). Additionally, there are significantly more homoeologs that show only some sharing of exon fractionation in comparisons of *Zea* genomes between different species and subspecies than in comparisons of accessions (Figure 3C).

To see if differences in conservation of homoeolog fractionation patterns may be related to constraint, we utilized a previously generated dataset of maize B73 genes for which dN/dS had been calculated for pairs of homoeologs (Yin et al. 2022). Of the 3,195 homoeologs that are in both our dataset and that of Yin et al. (2022), we find the distribution of dN/dS was similar between subgenomes, though M2 homoeologs have on average 0.01 higher dN/dS compared to M1 (p=0.048, Supp. Fig 9A). Further, we find that Zm B73 homoeologs that have some sharing of fractionation with another genome had higher dN/dS than the other categories (Sharing p<2e-16, Tukey p-values comparing “Frac.: Some Share” = 0 - 0.000005, Supp. Fig. 9B). When further separated into the types of genomes being compared with Zm B73, most were not statistically significant. The one exception was the M2 some sharing category in which comparisons with a *Tripsacum* genome had a significantly lower dN/dS than all *Zea* comparisons by 0.075-0.09 (Supp. Fig. 9C). To test if the distribution of Zm B73 dN/dS across sharing categories varied due to differences in the number of observations, we ran a permutation test to generate an expected 95% confidence interval based on sample size alone. Both subgenomes in the some shared category showed higher Zm B73 dN/dS than expected, suggesting that the higher dN/dS observed for this category is not due to differing sample sizes across categories (Supp. Fig. 9D). Thus, while genes aren’t neutral (dN/dS = 1) and thus under constraint (dN/dS < 1), some homoeologs appear to be under less constraint than others with significantly higher dN/dS.

### *Zea* genomes have more chromosomal rearrangements and descending dysploidy than *Tripsacum* genomes

While *Zea* genomes have long been known to have substantial chromosomal rearrangements relative to sorghum (Schnable et. al. 2011; Chia et. al. 2012; Lovell et. al. 2022), it was unclear if *Tripsacum* genomes shared this chromosomal history. We found all genomes retained two syntenic orthologous blocks relative to sorghum genome-wide (Figure 4A, Supp. Figure 4). Of the 18 modern chromosomes in *Tripsacum,* 14 comprise a full syntenic copy of a sorghum chromosome. The remaining 4 *Tripsacum* chromosomes either have one sorghum chromosome (Sb chromosomes 1 and 10) split across two modern *Tripsacum* chromosomes (Td chromosomes 4 and 10) or one sorghum chromosome inserted within another (Sb chromosome 5 inserted into Sb chromosome 2, creating Td chromosome 5) (Supp. Figure 10, Figure 4A). In contrast, *Zea* shows abundant chromosomal rearrangements such that 9 out of 10 modern *Zea* chromosomes are mosaics relative to sorghum chromosomes. A typical modern maize chromosome contains syntenic blocks to 2-4 sorghum chromosomes (Supp. Figure 10). Only *Zea* chromosome 7 shows synteny to a single sorghum chromosome (Figure 4A, Supp. Figure 10). Despite substantial rearrangements relative to sorghum, all *Zea* genomes have strong synteny to each other. The one exception is a known translocation from chromosome 10 to chromosome 9 in Zm Oh7b (Hufford et al., 2021). The same rearrangements in *Tripsacum* are also detected in *Zea*, suggesting those rearrangements occurred before the divergence of the two genera. Likewise, the numerous rearrangements shared across the *Zea* genomes likely happened before speciation in the *Zea* genus (Supp. Figure 4).

**Figure 4.**
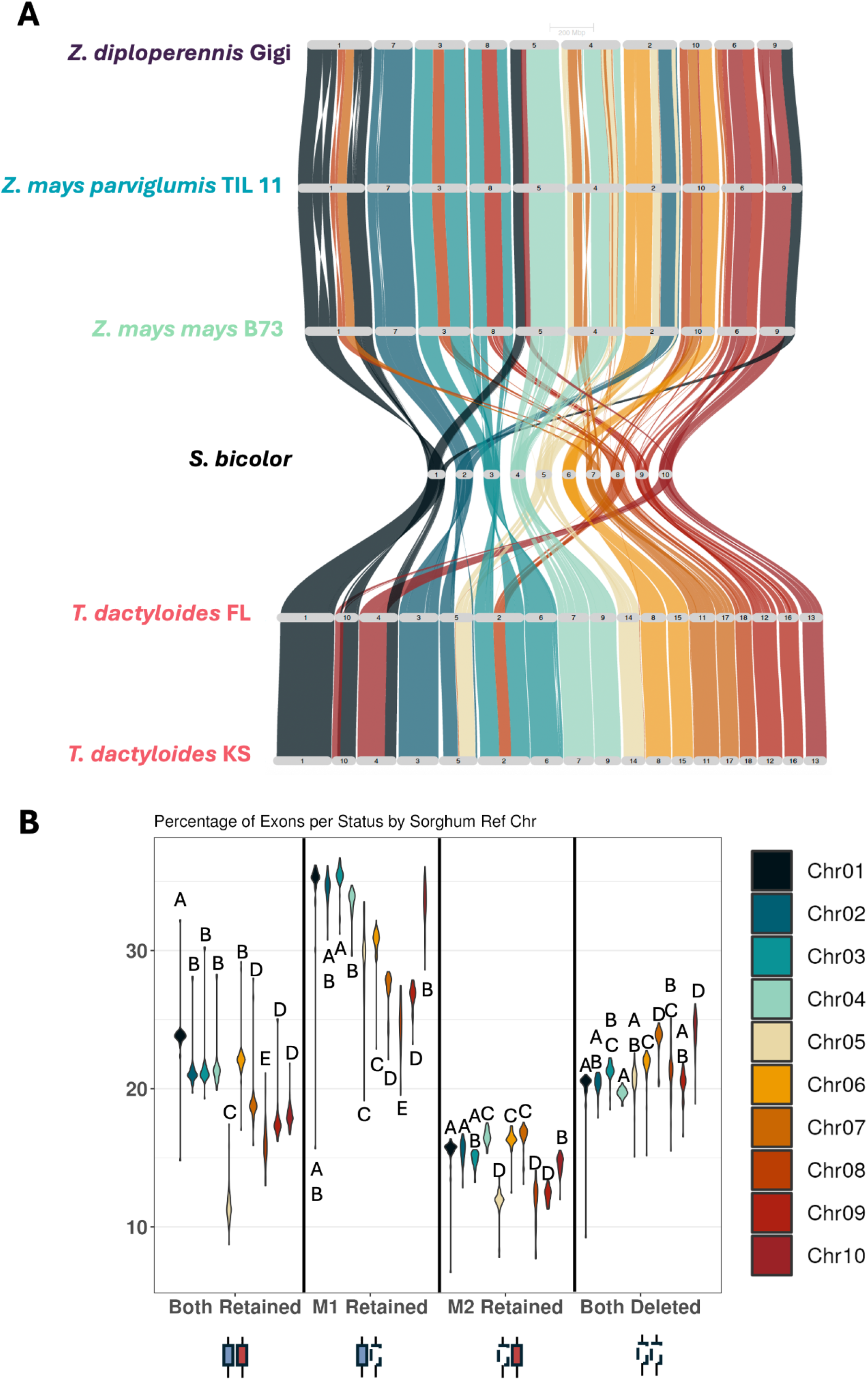
Spatial patterns of rearrangements and fractionation. (A) Riparian plot of a subset of representative genomes. Each chromosome is colored according to the ten sorghum chromosomes. Note each Tripsacinae genome matches 2:1 to each syntenic block in sorghum. (B) Percentage of homoeologous pairs of exons from each sorghum reference chromosome that show a given fractionation status. Differences in distributions across chromosomes were tested with ANOVA and post-hoc Tukey test for each fractionation category. Different letters indicate significant differences. Violin plots colored by sorghum reference chromosomes.

### Amount of fractionation varies by sorghum reference chromosome

Given that the polyploid ancestor from which all of these modern genomes are derived likely contained 20 chromosomes (Gaut & Doebley, 1997; Schnable et al., 2011) and the conserved synteny and collinearity between Andropogoneae genomes (Stitzer et al., 2025), we assume that the ancestral polyploid chromosome structure can be modeled by the modern assembly of sorghum in duplicate. Since the chromosomal rearrangement history differs greatly between *Tripsacum* and *Zea* and most fractionation differences also occur at the genus level, we investigated whether fractionation outcomes for homoeologous exon pairs differed based on ancestral starting chromosome, here assumed to be the sorghum chromosome. If fractionation occurred evenly across ancestral chromosomes, the proportion of exons in each fractionation category should be similar across sorghum chromosomes. However, we found significant differences across sorghum chromosomes (Figure 4B). In general, sorghum chromosomes that show a large number of rearrangements in the Tripsacinae (*e.g.*, sorghum chromosomes 5 and 7) or those that are smaller (*e.g.,* sorghum chromosomes 8 through 10) tend to show more fractionation of either both or 1 copy of an exon compared to sorghum chromosomes that have remained more intact or are larger (*e.g.*, sorghum chromosomes 1 through 3). For example, sorghum chromosome 1 has twice the average percentage of exons in the “both retained” category (mean =22.9%) as sorghum chromosome 5 (mean =11.4%). Syntenic regions of sorghum chromosome 5 show the greatest number of structural rearrangements in *Zea* across both subgenomes, where M1 is in at least 4 pieces and M2 is in 2 pieces (Figure 4A, Supp. Figure 10C,F). The M1 copy of sorghum chromosome 7, which is in three pieces across three *Zea* chromosomes and the smallest sorghum chromosome, chromosome 10, show the highest percentages of exons categorized as “both deleted” while sorghum chromosomes 1, 2, 3, 4, and 6 have the highest percentages of exons categorized as “both retained”. There are exceptions, such as the high percentage of M1 retained exons on sorghum chromosome 10. In general, these fractionation differences between ancestral chromosomes suggest there is a spatial component to the process of fractionation.

## Discussion

### Fractionation status is mostly shared, but differences are driven by higher retention of *Tripsacum* compared to *Zea*

Given that most fractionation is shared across all genomes or within genera, the majority of fractionation likely occurred either before the divergence of *Tripsacum* and *Zea* (∼5 MYA - 653 KYA, Schnable et al 2011, Chen et al 2022), or after the genera divergence but before *Zea mays* speciation and subspeciation events (∼653 KYA - 67 KYA, Chen et al 2022). Strong differences between genera were driven by higher retention in *Tripsacum* compared to *Zea* across multiple analyses. Across both subgenomes, ∼1% of exons show fractionation only in *Tripsacum* compared to ∼4% fractionated only across *Zea*. Across exons and genes, both copies are retained ∼5-8% more in *Tripsacum* than *Zea* genomes. Additionally, we found 20-35% of exon pairs show multiple fractionation statuses across genomes, which is greater than previous reports comparing maize accessions (∼2% Brohammer et al., 2018; ∼6% Hufford et al., 2021). This increase in observed fractionation differences between genomes is likely due to comparing more divergent lineages. This indicates several non-exclusive hypotheses. (1) Loss of an exon may be less deleterious than expected, particularly if a homoeologous exon remains intact; (2) there is selection for either retention or fractionation of a given gene or exon in a subset of lineages; (3) multiple mutational origins of exon loss may be common during fractionation.

It is well documented that polyploidy creates redundancy and buffers the effects of highly deleterious mutations, such as exon loss, so long as one homoeolog retains the original function (Cheng et al., 2018; Zhao et al., 2017). Previous studies show that more highly expressed genes are more likely to be retained in maize through purifying selection (Pophaly & Tellier, 2015; Renny-Byfield et al., 2017; Schnable et al., 2011). For a subset of maize B73 homoeologs with previous dN/dS estimates, both subgenome copies show signatures of purifying selection suggesting that these genes are under constraint, though M2 homoeologs dN/dS values were closer to 0 than their M1 counterparts. Despite this, ∼20% of exons per genome show both homoeologous exons have deletions, and ∼25% of the genes studied have deletions in every exon of both homoeologs. On top of the extensive occurrence of deletions, equal proportions of deletions in M1 sequences are expected to cause a frame-shift as opposed to maintenance of the reading frame. Frameshifting deletions occurred more often than non-frameshifting deletions for M2 homoeologs. Though the majority of deletions are small, previous work has demonstrated that fractionation in maize occurs through small initial deletions that grow into larger losses of sequence (Woodhouse et al., 2010). All together, this suggests that these deletions have widespread and likely loss-of-function effects on genes, particularly of the M2 subgenome. In cases where only parts of homoeologs were lost, other outcomes such as subfunctionalization, where a pair of homoeologs split the original function of the gene, and neo-functionalization, where a homoeolog gains a novel function, are possible (Bird et al., 2018; Cheng et al., 2018). Loss of exons or frameshifts could then be a source of adaptive diversity (Cheng et al., 2018). The members of the Tripsacinae subtribe, while centered in the neotropics, have dispersed to environments across extreme altitude, latitude, temperature, and precipitation gradients (Brink & De Wet, 1983; Hufford et al., 2012). Given the number of privately fractionated exons for individual wild species and subspecies, neo-functionalization and sub-functionalization might have supported adaptation to these different environments, though this would require experimental validation to prove. Thus, the frequency at which both homoeologs were completely lost and the possibility of adaptive potential in altered gene structures suggests that perhaps exon loss may not be as deleterious post-polyploidy.

Including a broader sampling of lineages from across the subtribe allowed us to detect fractionation shared by non-monophyletic lineages, approximately 10% of all exons. Such discordance could be the product of multiple mutational origins converging on exon loss or incomplete lineage sorting. Based on estimates of expected ILS, we expected discordance to be more commonly observed between less diverged lineages. We set these expectations using an equation for neutral variants in simplified three lineage scenarios (see methods and Supp. Figure 6). We expect these estimates to represent an upper bound of possible ILS as more complicated patterns of discordance are less likely than these simple three lineage patterns. Likewise, deletion variants are unlikely to be neutral, which should result in lower than expected levels of discordance. Finally, fractionation calls per exon represent a collapsing of complex deletion genotypes to create a binary variable of fractionation. This collapsing obscures the presence of multiple deletions within a single exon and increases the likelihood of calling independent exon loss by different deletions “shared”, and thus creating homoplasy. In comparing our expectations to observations, we found the most diverged lineages showing the greater overall discordance, opposite our expectations. Though we predict ILS should be commonly observed, this reversal of expectations suggests multiple mutations resulting in convergent exon loss likely play a larger role in this observed discordance, particularly discordance between *Zea* and *Tripsacum* lineages. At the gene level, few instances of completely different fractionation patterns were observed, but partial differences in comparisons of a *Tripsacum* genome to any *Zea* genome were more common than differences between *Zea* genomes. Given that these are the most divergent genome comparisons, future work could dissect whether these differences stem from a shared deletion in the common ancestor expanding in one lineage and not the other or if deletions in the same region occurred independently.

Overall, our findings agrees with previous work that fractionation began soon after the WGD event (Li et al., 2016; Sankoff et al., 2010; Schnable et al., 2011), and that differences have accumulated between sister lineages (Brohammer et al 2018, Hufford et al 2021).

### Sharing of gene loss likely driven by relaxed purifying selection

In comparisons of fractionated genes in two genomes, on average 75% of genes were fractionated in the exact same way. However, that number drops to closer to 60% when comparing a *Tripsacum* and *Zea* genome to each other, suggesting that differences in which exons are deleted are correlated with divergence between two genomes. If multiple mutational origins underlie fractionation of a given homoeolog, this could either be due to relaxed purifying selection allowing multiple deletions to accumulate or selection for loss leading to convergent fractionation. We sought to tease these two possibilities apart using a previously published dataset (Yin et al., 2022). For the subset of 3,195 genes which Yin et al. (2022) reported dN/dS in maize accession B73, M2 homeologs overall had a slightly higher dN/dS than M1 homoeologs and genes that had some sharing, but not complete sharing, of fractionation between genomes had higher dN/dS than other sharing categories. While all dN/dS average estimates were less than 1 and thus indicate constraint, the higher dN/dS of these subsets suggest constraint is weaker on M2 homoeologs and the subset of Zm B73 homoeologs with some fractionation sharing with other genomes. Relaxed constraint on M2 fits with the biased fractionation of M2 over M1 homoeologs, where M1 homoeologs are likely retaining original functions while M2 accumulate deletions, as has been described in previous maize studies (Renny-Byfield et al, 2017, Schnable et al., 2011). The higher dN/dS of M2 homoeologs in the some shared fractionation category between Zm B73 and other *Zea* genomes versus Zm B73 and *Tripsacum* genomes could be due to differences in the homoeologs observed in those categories.

It is worth noting that this dN/dS dataset represents roughly one quarter of all the genes investigated in our study, and may not be a completely representative sample as Yin et al. (2022) include only homoeologs for which sequence can be found for both subgenomes. Thus, genes that are retained as singletons or have been fractionated in both subgenomes are not included, which may have different dN/dS distributions. Calculating dN/dS for these categories may be difficult as interpreting large proportions of indels in dN/dS calculations can be challenging, and in our study, on average 35% of gene coding sequence has been lost from each homoeolog. Likewise, these are only the dN/dS values reported for one maize accession, B73, relative to sorghum. Distributions of dN/dS may vary for the same genes across other genomes in ways we cannot detect with this dataset. Based on previous work, we expect that the majority of fractionated genes would have higher dN/dS than retained genes as fractionated genes most often face gene death (Li et al., 2021, Wendel 2015).

Overall, the higher Zm B73 dN/dS of homoeologs that show some sharing of fractionation versus complete or no sharing with other genomes suggests that those homoeologs are under less constraint, and thus variable fractionation of a homoeolog across genomes may be due to relaxed purifying selection rather than convergent positive selection.

### Differences in exon retention between genera may be related to separate chromosomal rearrangement histories

In all of our analyses, differences were greatest and often only statistically significant between genera than within a genus (*i.e., Tripsacum* vs. *Zea* rather than *Zea* vs. *Zea*). *Tripsacum* had significantly higher retention of homoeologous pairs and fewer exons lost per gene than *Zea.* Given our expectation that the process and outcomes of fractionation would be similar across lineages descending from the same polyploid ancestor, we considered what may underlie these differences. A distinction between the genera is the significant difference in chromosomal rearrangements relative to sorghum and the current haploid chromosome number. *Tripsacum* has fewer rearrangements and 18 smaller chromosomes while *Zea* contains many rearrangements and 10 larger chromosomes (Maguire 1961). Descending dysploidy, or a decrease in chromosome number, via translocations can occur when two non-homologous chromosomes have double stranded breaks which are resolved through nonhomologous end joining across chromosomes or non-allelic homologous recombination between repetitive sequences found on both chromosomes (Lysák and Schubert 2013). The extensive rearrangements of *Zea* chromosomes occurs relatively close to the WGD given the strong synteny of the genus. While the mechanism giving rise to this rapid karyotypic change is unidentified, heterochromatic knob sequence is enriched near several breakpoints (Stitzer et al., 2025). Knob tandem repeats mediate meiotic drive in maize, and the genes responsible for drive expand at a similar time to the divergence of the genera *Zea* and *Tripsacum* (Rhoades 1942, Dawe et al 2018, Chen et al 2022).

Grasses show limited descending dysploidy, with few reductions in chromosome number post-polyploidy (Stitzer et al., 2025, Ma et al., 2024, Murat et al., 2014, Salse 2016, Scarlett et al., 2023). A broader taxonomic view shows extensive chromosomal rearrangements are common after polyploidy (Carta et al., 2020, Li et al., 2021, Salse 2012), with resynthesis experiments of *Brassica napus* showing chromosomal rearrangements occurring as early as the initial meiosis generation and becoming fixed after as few as 3 or 5 generations of selfing (Szadkowski et al., 2010). These differences between monocots and eudicots indicate potential lineage specificity in chromosomal structural rearrangement post-polyploidy. Furthermore, recent studies of *Microlepidieae*, a crucifer tribe endemic to Australia and New Zealand, have likewise shown increased fractionation with greater numbers of chromosomal rearrangements and chromosome number reductions (Mandáková et al., 2017b). Changes in life history (*e.g.*, transition from perennialism to annualism) could increase recombination rates, chromosomal rearrangements, and/or fractionation rates; such trends were observed, though not statistically significant, in the *Microlepidieae* (Mandáková et al., 2017b). These trends also fit with our observations of perennial *Tripsacum* with fewer rearrangements and less fractionation vs. largely annual *Zea* species with more rearrangements and more fractionation. The transition between perennialism and annualism could also result in different generation times as annuals generally have more generations per unit time than perennials and has been linked to higher rates of molecular evolution in some cases (see Friedman, 2020 for a more comprehensive review), which could affect observed differences in fractionation. Given that there is only one perennial to annual transition within these lineages, it is difficult to disentangle differing numbers of generations with the observed differences in fractionation extent. Notably, *Zea diploperennis* is perennial and yet shows a similar extent of fractionation as seen in the annual *Zea* species. Lower effective population sizes (Ne) are also related to higher fixation of chromosomal rearrangements (Lande 1979). Given the recent release of these genomes, the only estimates of Ne currently published are for *Zea* species over the recent 100KY (Chen et al., 2022), far more recent than the divergence from *Tripsacum* at 650 KYA. Thus it remains to be seen how estimates of population size differ between these genera. The causal factors leading to either chromosomal rearrangement or chromosomal conservation post-polyploidy and their relationship to fractionation outcomes remain an open question. However, our observations that fractionation outcomes are non-random with respect to their starting chromosomal location and the larger extent of sequence loss in genomes with more rearrangements suggests a spatial component to the process of fractionation.

## Conclusion

The larger patterns of biased subgenome fractionation occurring shortly after the polyploidy event observed from investigations in maize also hold true in this expanded study that includes wild species across the Tripsacinae. However, there are strong differences in the amount of fractionation, the way homoeologs are fractionated, and the amount of fractionation relative to the ancestral chromosomes between *Zea* and *Tripsacum*. This suggests that fractionation followed two different trajectories within the subtribe, perhaps related to the differences in chromosomal rearrangements which characterize the genera. All together, this indicates that the process of fractionation is more dynamic in the Tripsacinae than expected, and that while reduced gene content is a highly repeated outcome of rediploidization across Angiosperms, how and what sequences are lost may differ between lineages in ways previously not anticipated.

## Materials and Methods

### Data

The publicly available pseudomolecule assemblies for two *Zea mays* ssp. *mexicana* accessions (Zx: TIL25, TIL18), two *Zea mays* ssp*. parviglumis* accessions (Zv: TIL01, TIL11), one *Zea mays* ssp. *huehuetenagensis* accession (Zh: RIMHU001), two *Zea diploperennis* accessions (Zd: Gigi, Momo), one *Zea nicaraguensis* accession (Zn: PI615697), two *Tripsacum dactyloides* accessions (Td: FL, KS), and one *Anatherum virginicum* (Av) accession (Stitzer et al., 2025) were used in conjunction with the existing NAM *Zea mays* ssp. *mays* genomes (26 accessions, Zm: Tropical: Tzi8, NC358, NC350, Ki3, Ki11, CML69, CML52, CML333, CML322, CML277, CML247, CML228, CML103. Mixed: Tx303, Mo18W, M37W. Temperate: P39, HP301, Oh7b, Oh43, MS71, M162W, Ky21, IL14H, B97, B73). *Sorghum bicolor* 313 (sorghum) (McCormick et al., 2017) and *Anatherum virginicum* (Stitzer et al., 2025) from the subtribes Andropogoninae and Sorghinae respectively (Welker et al. 2020) diverged from each other approximately 10.6 MYA and are similar phylogenetic distances from the Tripsacinae (approximately 18.9 MYA)(Stitzer et al., 2025). We use these modern species’ genomes as models for the extinct, ancestral diploid genomes which hybridized during the Tripsacinae WGD event between 5 and 12 MYA. No closer relatives could be used as an outgroup to this subtribe.

### Whole genome alignments and subgenome assignments relative to sorghum

Based on segment length and number of aligned segments, we filtered syntenic orthologous regions identified by GENESPACE (Lovell et al., 2022) that did not match the expected 2:1 relationship arising from the Tripsacinae WGD. Small regions around 1MB in length and deviating from this expectation (3 or more to 1) were removed as likely remnants from an older WGD that is shared by all members of the *Poaceae*, which includes the Tripsacinae and sorghum (Paterson et al., 2004). Given the strong synteny across *Zea*, we manually assigned subgenomes to individual *Zea* orthologous syntenic regions based on synteny with sorghum and the subgenome assignments from Schnable et al. (2011). We used the syntenic regions identified by GENESPACE (Lovell et. al., 2022) between each *Tripsacum* genome and Zm B73 to manually assign subgenome identity from Schnable et al (2011). For example: *Tripsacum* chr13 is syntenic to sorghum Chr10 and Zm B73 chr6. Given the region of B73 chr6 syntenic to sorghum Chr10 is assigned the M2 subgenome, *Tripsacum* chr13 is assigned to M2. The same process was used to resolve the subgenome assignment of the translocation in Zm Oh7b.

Each Tripsacinae genome was aligned to sorghum using AnchorWave (Song et al., 2022) with sorghum as reference and parameters -R 2 and -Q 1. For *Z. nicaraguensis* and *Z. diploperennis* assemblies which had low mapping rates, likely due to assembly artifacts of highly heterozygous individuals, we made additional alignments with -R 4 and -Q 1 (labeled 4to1). Resulting MAFs were split first by Tripsacinae sequence name and then by the sorghum sequence name using mafSplit (Perez et al., 2024). Using subgenome identity assigned to each Tripsacinae-sorghum chromosome pair, the split MAFs were concatenated by subgenomes into “Bin1” and “Bin2” MAF files corresponding to the M1 and M2 subgenomes, respectively. Concatenated subgenome MAFs were converted to GVCF using the MAFToGVCF phg plugin (Bradbury et al., 2022). GVCFs’ alignment quality was evaluated by VCFMetricsPlugin (Bradbury et al 2007).

To remove any potential lineage-specific sorghum exons, we identified a set of conserved gene models to use for all downstream analyses. This gene set was identified through reciprocal liftover of sorghum 313 and *A. virginicum* annotations using Liftoff (Shumate and Salzberg 2021). Sorghum and *A. virginicum* annotations with the exact same exon structures (exact same exon number and exon lengths) were kept as outgroup reference exons (12,157 genes, 69,269 exons). Given the potential for different definitions of exons, CDS, and UTR regions across gffs, we verified that these reference sequences in sorghum did not overlap with features annotated as UTRs.

### Calling and characterizing deletions

For each sorghum chromosome, M1 and M2 GVCFs were combined across all genomes except *Z. nicaraguensis* to create a VCF using GATK GenomicsDBImport and GenotypeGVCFs tools (Van der Auwera and O’Connor 2020). VCFs were filtered for indel variants using GATK SelectVariants (Van der Auwera and O’Connor 2020) and reformatted using bcftools annotate (Danecek et al., 2021). Custom scripts further filtered insertion variants and converted the VCF to bed format. Lengths of deletions were calculated as the length of the REF allele minus the length of the ALT allele to create end coordinates and also maintain strandedness. Further filtering of deletions left only those intersecting reference exons as “exonic deletions” using bedtools intersect (Quinlan and Hall 2010). Deletions that reciprocally overlap at least 80% of their sequence were collapsed. New collapsed genotypes for reference correspond to only “0” genotypes, alternative if at least one deletion was genotyped “1”, and NA or “.” if no call was made. Deletions were considered frame shifting if the deletion length was not a multiple of 3.

### Calling fractionation status

AnchorWave (Song et al., 2022) guides alignment by identifying syntenic blocks, and thus regions beyond syntenic blocks may not be aligned and thus not included in the GVCF. To identify reference exons that were aligned, each GVCF was converted to a bed file by a custom script and then intersected with the sorghum reference exon bedfile using bedtools2 (Quinlan and Hall 2010). Reference exons within the GVCF-derived bed file were classified as “aligned” and all others as “unaligned”. All genomes were checked to make sure that the number of aligned and unaligned exons summed to the total 69,269 reference exons.

Each GVCF-derived bed file was filtered for deletions, defined as the length of the reference allele being longer than the length of an alternative allele that was not <NON_REF>. Deletion lengths were assigned as the length of the reference allele minus the length of the alternative allele. These deletions’ coordinates were intersected with the aligned exons for that GVCF (Quinlan and Hall 2010). If the aligned reference exon intersected with at least one deletion, that exon was called “fractionated”. Otherwise, the exon was called “retained” (Supp. Figure 1). Each set of retained and fractionated calls by genome and subgenome was joined by CDS ID using a custom R script (“fractionation-calling.R”) to create a dataframe where each row indicated a reference exon, each subgenome of each query genome was a column, and the value within each cell referred to fractionation status (0 = retained, 1 = fractionated, NA = unaligned)(Supp. Data 3).

For analyses summarizing fractionation of an entire gene, multiple thresholds were used. The broadest was at least one exon within the gene called fractionated (“at least one” or “only1”) and the strictest required all exons to be fractionated (“all”). Between these two cutoffs were at least ⅓ or ½ of the exons called fractionated (“⅓” or “third” and “½” or “half” respectively). For all cutoffs, if a gene did not meet the cutoff for fractionation calling, it was considered retained (Supp. Data 4). To estimate the amount of a homoeolog being fractionated, we calculated the number of exons called fractionated out of the total exons within the homoeolog as well as the percentage of base pairs overlapping a deletion out of the total base pairs in the coding sequence of a gene model.

There was a significantly larger percentage of unaligned exons within *Z. nicaraguensis* and *Z. diploperennis* compared to other Tripsacinae genomes. These genomes were assembled from heterozygous individuals and contain multiple haplotypes. Increasing the alignment ratio from 2:1 to 4:1 aligned thousands more exons for each *Z. diploperennis* genomes (Zd Gigi = 2933, ZdMomo = 5241), but only 341 more exons for *Z. nicaraguensis.* This is likely due to the more fragmented *Z. nicaraguensis* genome assembly rather than biological differences. Given that *Z. diploperennis* genomes had better alignments overall that were substantially improved compared to *Z. nicaraguensis*, the “4to1” *Z. diploperennis* alignments were used in all downstream analyses, but *Z. nicaraguensis* was removed.

All scripts for this analysis can be found at: https://github.com/Snodgras/Zea_Fractionation

### Sharing of fractionation

Pairs of homoeologous exons and genes were categorized by retention, loss, and alignment status by subgenome and genome, and then summary statistics were calculated. To relate the sharing of fractionation to divergence time between genomes, we use a phylogeny of the query genomes created from previous phylogenies (Stitzer et al., 2025, Hufford et al., 2012; Hufford et al., 2021, Chen et al., 2022). In calculating the percentage of exons fractionated along a given clade, only exons that were aligned across all genomes were included. Exons that exhibited non-monophyletic patterns were not assigned a branch.

To generate an upper bound expectation for the amount of discordance or non-monophyly given incomplete lineage sorting (ILS), we use a simplified phylogeny and consider only 3 lineages at a time for each type of discordance or ILS case we test (Supp. Figure 6). Estimates of divergence times and Ne come from Chen et al. (2022). We estimate the expected probability of ILS using equation 6.1 from Coop’s open source population genetics notes (https://github.com/cooplab/popgen-notes). To count observations of discordance, one genome is randomly selected to represent the lineage, and counts of exons showing each type of discordance is recorded. Under each scenario, discordance means either one of the sister lineages and the outlier lineage shows fractionation of an exon while the remaining sister lineage retains the exon. Sampling of genomes is repeated 1000 times to create a distribution. Both expected and observed counts were converted to percentage of total aligned exons per subgenome (M1 = 59,164, M2 = 44,321) for easier comparison across figures. Since we only consider simple, three lineage cases of discordance and reduce all accessions down to a single lineage, the total percentages of observed discordance are lower (approximately 2%) than when all genomes and more complicated patterns of discordance are observed (approximately 10%).

To categorize differential fractionation across genomes, we counted the number of observed aligned statuses (“both retained”, “M1 only”, “M2 only”, “both lost”) of each exon. Here we did not require that all exons pairs be aligned to be included, so the denominator is the total number of exons (69,269). This results in slightly different percentages compared to the phylogeny related analysis (65% complete sharing versus 80% complete sharing).

### Gene Expression Analysis

In order to evaluate gene expression, sorghum gene coordinates were lifted over onto query genome coordinates. We did this by converting each MAF to chain files and then lifting over coordinates from sorghum to the query genome coordinate system using Crossmap (Zhao et al., 2014). Given limited gene expression data available for the wild accessions of our study, we evaluated gene expression and fractionation using publicly available leaf tissue transcripts for the 26 maize accessions. Bam files were downloaded from maizeGDB (Woodhouse et al., 2021). Based on the position of fractionated and retained genes in each of the NAM lines, we prepared a gff file. We use htseq-count (Putri et al., 2022) to calculate the number of reads for each fractionated and retained gene. Given that not all sorghum reference genes could be mapped onto NAM coordinates because they were completely deleted, we normalized read counts using a slightly modified TPM, such that reads were normalized using ((number of reads mapped to a gene / length of that gene) / number of reads mapped to all lifted over genes in that NAM line) * 1000000.

### Gene Ontology Analysis

Gene ontology enrichment analysis was performed using the PANTHER database tools (Thomas et al., 2022). We used the PANTHER GO-Slim categories as they are more manually curated and of higher confidence than the broader GO categories for the three domains: biological, cellular, and molecular processes. We used our set of ∼12K highly conserved genes as the reference list instead of all the genes within the reference sorghum gene annotation. Fisher’s exact test with false discovery rate p-value adjustments were used for all tests. All GO analyses were performed solely using sorghum genes and not any Tripsacinae annotations.

### Convergence of fractionation by gene

To evaluate potential convergent loss of the same homoeologs, we conducted all pair-wise comparisons between genomes for each individual homoeolog. Homoeologs where any exon was unaligned were ignored for this analysis. If all of the exons that were fractionated in each genome were the same, that homoeolog was considered “complete share”. If both genomes show fractionation but none of the fractionated exons were the same, the homoeolog was considered “no share”. If all exons were shared or different due to both or one genome fully retaining a homoeolog respectively, those were categorized separately to parse similarities and differences caused by retention from those solely by fractionation. If some exons shared fractionation status, but not all, the homoeolog was considered “some shared”.

### dN/dS Analysis

We intersect Zm B73 M1 and M2 dN/dS measures relative to sorghum from Yin et al. (2022) and the sorghum gene models from our study to generate a subset of 3,195 genes with Zm B73 M1 and M2 dN/dS. For each of these 3,195 genes, we observe if Zm B73 shared fractionation with each of the other genomes, such that if both genomes have a fractionated homoeolog that is fractionated in the exact same way, it is “complete share”, fractionated in overlapping ways, “some share”, or in different, non-overlapping ways, “no share”. If one genome fully retains the homoeolog while the other genome shows fractionation, it is “Retained: No Share”. B73 dN/dS is then compared across categories, always keeping M1 and M2 values separate. The overall distribution of B73 dN/dS between subgenomes was tested using paired t-test, pairing homoeologs between the subgenomes. All other comparisons were tested using ANOVA and post-hoc Tukey tests. Given unequal occurrences across the sharing categories, with more instances of “complete share” than “some share” than “no share”, we used a permutation test to see if our observations could be due to unequal sample sizes. We randomly reassigned dN/dS values to subgenome and sharing categories and recorded the average dN/dS for each category 1000 times. 95% confidence intervals were calculated from these distributions and compared to the observed average dN/dS per category.

### Spatial organization of fractionation

For each sorghum reference chromosome, the counts of homoeologous exon pairs in each fractionation status category (“Both Retained”, “Both Deleted”, “M1 Retained”, “M2 Retained”) was calculated. Percentages were calculated by count of exons within a category divided by the total number of exons on a given sorghum reference chromosome. The significance of differences in percent exons in each category across sorghum reference chromosomes was evaluated using an ANOVA.

## Acknowledgements

We would like to acknowledge our funding sources: DGE#1744592, NSF-IOS 1822330, and NSF IOS 2305694, MRI1726447, and MRI2018594, which supported the sequencing, computational, and labor costs of this project. We thank the practical haplotype graph team, especially Ana Helene Berthel and Zach Miller, who generously helped troubleshoot the MAF to GVCF plugin in Tassel5. We thank Jonathan Wendel for his advice on this project and general enthusiasm and encouragement in all things polyploid. We thank Elli Cryan for her insights on dN/dS pipelines. We thank Jeff Ross-Ibarra and Graham Coop for their feedback on ideas, methods, and general willingness to be sounding boards, particularly with regards to ILS. This research was also supported by the U.S. Department of Agriculture, Agricultural Research Service, Project Number [5030-21000-072-00-D] through the Corn Insects and Crop Genetics Research Unit in Ames, Iowa. Mention of trade names or commercial products in this publication is solely for the purpose of providing specific information and does not imply recommendation or endorsement by the U.S. Department of Agriculture. USDA is an equal opportunity provider and Employer. This research used resources provided by the SCINet project and/or the AI Center of Excellence of the USDA Agricultural Research Service, ARS project numbers 0201-88888-003-000D and 0201-88888-002-000D.

## Data Availability Statement

All scripts underlying this article are available on github at https://github.com/Snodgras/Zea_Fractionation. The genome assemblies and annotations underlying this article are available at maizeGDB (Woodhouse et al., 2021, https://doi.org/10.1186/s12870-021-03173-5). Exons used for the high quality set are available in supplemental file ref_Sb313_cds.bed (Supp. Data 2). Estimates of dN/dS are available in the Yin et al (2022) and in its online supplementary material. Fractionation calls at the exon and gene level are available in Supp. Data 3 and 4 respectively.

## Supplemental Files

Supplemental Data 1: GO Terms

Supplemental Data 2: ref_Sb313_cds.bed

Supplemental Data 3: full.fractionation.status.SuppData3.tsv

Supplemental Data 4: gene.fractionation.SuppData4.tsv

## Supplemental Tables

**Supp. Table 1.** Counts of reference exons of each fractionation status across all genomes. Included are the counts for *Zea diploperennis* with both standard 2:1 and 4:1 alignments to Sorghum. The total number of reference exons is 69,269.

| Genome | Both Retained | M1 Retained | M2 Retained | Both Lost | M1 Retained, M2 NA | M1 NA, M2 Retained | M1 Lost, M2 NA | M1 NA, M2 Lost | Both NA |
| --- | --- | --- | --- | --- | --- | --- | --- | --- | --- |
| Zm B73 | 14104 | 22862 | 10614 | 14807 | 2994 | 717 | 1897 | 859 | 415 |
| Zm B97 | 14142 | 22624 | 10380 | 14655 | 3278 | 763 | 2031 | 1001 | 395 |
| Zm IL14H | 14203 | 22639 | 10483 | 14714 | 3173 | 797 | 1892 | 1014 | 354 |
| Zm Ky21 | 14303 | 22768 | 10710 | 14667 | 3033 | 582 | 1939 | 917 | 350 |
| Zm M162W | 14210 | 22693 | 10586 | 14727 | 3261 | 609 | 1903 | 977 | 303 |
| Zm MS71 | 14031 | 22761 | 10875 | 14946 | 2969 | 633 | 1831 | 870 | 353 |
| Zm Oh43 | 14227 | 22472 | 10447 | 14729 | 3233 | 735 | 2043 | 952 | 431 |
| Zm Oh7b | 14054 | 22701 | 10378 | 14720 | 3024 | 930 | 1934 | 1088 | 440 |
| Zm P39 | 14189 | 22569 | 10455 | 14857 | 3105 | 781 | 1930 | 940 | 443 |
| Zm HP301 | 14238 | 22797 | 10368 | 14823 | 2918 | 862 | 1834 | 1087 | 342 |
| Zm M37W | 14231 | 22683 | 10517 | 14740 | 3102 | 737 | 1902 | 981 | 376 |
| Zm Mo18W | 14207 | 22570 | 10616 | 14687 | 3241 | 672 | 1954 | 909 | 413 |
| Zm Tx303 | 14065 | 22957 | 10625 | 14653 | 2973 | 709 | 1978 | 923 | 386 |
| Zm CML103 | 14350 | 22831 | 10576 | 14814 | 2871 | 648 | 1834 | 946 | 399 |
| Zm CML228 | 13968 | 22318 | 10348 | 14223 | 3443 | 869 | 2302 | 1336 | 462 |
| Zm CML247 | 14228 | 22674 | 10573 | 14870 | 3076 | 670 | 1856 | 929 | 393 |
| Zm CML277 | 14206 | 22804 | 10479 | 14725 | 3179 | 674 | 1901 | 921 | 380 |
| Zm CML322 | 14195 | 22895 | 10369 | 14905 | 2927 | 730 | 1960 | 943 | 345 |
| Zm CML333 | 14227 | 22789 | 10652 | 14787 | 3095 | 642 | 1868 | 875 | 334 |
| Zm CML52 | 14014 | 22557 | 10546 | 14660 | 3300 | 746 | 1974 | 967 | 505 |
| Zm CML69 | 14172 | 22764 | 10607 | 14823 | 3259 | 584 | 1880 | 798 | 382 |
| Zm Ki11 | 14268 | 22957 | 10481 | 14746 | 2844 | 730 | 1717 | 981 | 545 |
| Zm Ki3 | 14281 | 22970 | 10459 | 14973 | 2899 | 667 | 1745 | 915 | 360 |
| Zm NC350 | 14146 | 22789 | 10528 | 14672 | 3024 | 704 | 1999 | 946 | 461 |
| Zm NC358 | 14074 | 22521 | 10655 | 14674 | 3332 | 651 | 2026 | 1000 | 336 |
| Zm Tzi8 | 14201 | 22640 | 10584 | 14615 | 3240 | 602 | 2093 | 940 | 354 |
| Zv TIL11 | 14068 | 22672 | 10542 | 14743 | 3206 | 749 | 1963 | 824 | 502 |
| Zv TIL01 | 14220 | 22600 | 10373 | 14727 | 3219 | 743 | 1963 | 991 | 433 |
| Zx TIL18 | 14219 | 22712 | 10629 | 14586 | 3142 | 709 | 1925 | 944 | 403 |
| Zx TIL25 | 13979 | 22526 | 10723 | 14540 | 3285 | 702 | 2184 | 903 | 427 |
| Zh RIMHU001 | 13916 | 22084 | 10267 | 14207 | 3519 | 906 | 2414 | 1202 | 754 |
| Zd Gigi | 13042 | 20019 | 9648 | 12593 | 5528 | 1716 | 3473 | 2245 | 1005 |
| Zd Momo | 12088 | 18820 | 8800 | 12044 | 7244 | 1943 | 4509 | 2778 | 1043 |
| Zd Gigi_4to1 | 13746 | 20935 | 10135 | 13270 | 4554 | 1306 | 2698 | 1769 | 856 |
| Zd Momo_4to1 | 13290 | 20588 | 9741 | 13217 | 5283 | 1249 | 3083 | 1932 | 886 |
| Td FL | 19267 | 20703 | 9415 | 13932 | 2387 | 701 | 1523 | 797 | 544 |
| Td KS | 16735 | 18596 | 8392 | 12497 | 7027 | 679 | 3853 | 878 | 612 |

**Supp. Table 2.** Percentages of reference exons of each fractionation status across all genomes. Included are the percentages for *Zea diploperennis* with both standard 2:1 and 4:1 alignments to Sorghum. Rounded to 2 decimal places.

| Genome | Both Retained | M1 Retained | M2 Retained | Both Lost | M1 Retained, M2 NA | M1 NA, M2 Retained | M1 Lost, M2 NA | M1 NA, M2 Lost | Both NA |
| --- | --- | --- | --- | --- | --- | --- | --- | --- | --- |
| Zm B73 | 20.36% | 33.00% | 15.32% | 21.38% | 4.32% | 1.04% | 2.74% | 1.24% | 0.60% |
| Zm B97 | 20.42% | 32.66% | 14.99% | 21.16% | 4.73% | 1.10% | 2.93% | 1.45% | 0.57% |
| Zm IL14H | 20.50% | 32.68% | 15.13% | 21.24% | 4.58% | 1.15% | 2.73% | 1.46% | 0.51% |
| Zm Ky21 | 20.65% | 32.87% | 15.46% | 21.17% | 4.38% | 0.84% | 2.80% | 1.32% | 0.51% |
| Zm M162W | 20.51% | 32.76% | 15.28% | 21.26% | 4.71% | 0.88% | 2.75% | 1.41% | 0.44% |
| Zm MS71 | 20.26% | 32.86% | 15.70% | 21.58% | 4.29% | 0.91% | 2.64% | 1.26% | 0.51% |
| Zm Oh43 | 20.54% | 32.44% | 15.08% | 21.26% | 4.67% | 1.06% | 2.95% | 1.37% | 0.62% |
| Zm Oh7b | 20.29% | 32.77% | 14.98% | 21.25% | 4.37% | 1.34% | 2.79% | 1.57% | 0.64% |
| Zm P39 | 20.48% | 32.58% | 15.09% | 21.45% | 4.48% | 1.13% | 2.79% | 1.36% | 0.64% |
| Zm HP301 | 20.55% | 32.91% | 14.97% | 21.40% | 4.21% | 1.24% | 2.65% | 1.57% | 0.49% |
| Zm M37W | 20.54% | 32.75% | 15.18% | 21.28% | 4.48% | 1.06% | 2.75% | 1.42% | 0.54% |
| Zm Mo18W | 20.51% | 32.58% | 15.33% | 21.20% | 4.68% | 0.97% | 2.82% | 1.31% | 0.60% |
| Zm Tx303 | 20.30% | 33.14% | 15.34% | 21.15% | 4.29% | 1.02% | 2.86% | 1.33% | 0.56% |
| Zm CML103 | 20.72% | 32.96% | 15.27% | 21.39% | 4.14% | 0.94% | 2.65% | 1.37% | 0.58% |
| Zm CML228 | 20.16% | 32.22% | 14.94% | 20.53% | 4.97% | 1.25% | 3.32% | 1.93% | 0.67% |
| Zm CML247 | 20.54% | 32.73% | 15.26% | 21.47% | 4.44% | 0.97% | 2.68% | 1.34% | 0.57% |
| Zm CML277 | 20.51% | 32.92% | 15.13% | 21.26% | 4.59% | 0.97% | 2.74% | 1.33% | 0.55% |
| Zm CML322 | 20.49% | 33.05% | 14.97% | 21.52% | 4.23% | 1.05% | 2.83% | 1.36% | 0.50% |
| Zm CML333 | 20.54% | 32.90% | 15.38% | 21.35% | 4.47% | 0.93% | 2.70% | 1.26% | 0.48% |
| Zm CML52 | 20.23% | 32.56% | 15.22% | 21.16% | 4.76% | 1.08% | 2.85% | 1.40% | 0.73% |
| Zm CML69 | 20.46% | 32.86% | 15.31% | 21.40% | 4.70% | 0.84% | 2.71% | 1.15% | 0.55% |
| Zm Ki11 | 20.60% | 33.14% | 15.13% | 21.29% | 4.11% | 1.05% | 2.48% | 1.42% | 0.79% |
| Zm Ki3 | 20.62% | 33.16% | 15.10% | 21.62% | 4.19% | 0.96% | 2.52% | 1.32% | 0.52% |
| Zm NC350 | 20.42% | 32.90% | 15.20% | 21.18% | 4.37% | 1.02% | 2.89% | 1.37% | 0.67% |
| Zm NC358 | 20.32% | 32.51% | 15.38% | 21.18% | 4.81% | 0.94% | 2.92% | 1.44% | 0.49% |
| Zm Tzi8 | 20.50% | 32.68% | 15.28% | 21.10% | 4.68% | 0.87% | 3.02% | 1.36% | 0.51% |
| Zv TIL11 | 20.31% | 32.73% | 15.22% | 21.28% | 4.63% | 1.08% | 2.83% | 1.19% | 0.72% |
| Zv TIL01 | 20.53% | 32.63% | 14.97% | 21.26% | 4.65% | 1.07% | 2.83% | 1.43% | 0.63% |
| Zx TIL18 | 20.53% | 32.79% | 15.34% | 21.06% | 4.54% | 1.02% | 2.78% | 1.36% | 0.58% |
| Zx TIL25 | 20.18% | 32.52% | 15.48% | 20.99% | 4.74% | 1.01% | 3.15% | 1.30% | 0.62% |
| Zh RIMHU001 | 20.09% | 31.88% | 14.82% | 20.51% | 5.08% | 1.31% | 3.48% | 1.74% | 1.09% |
| Zd Gigi | 18.83% | 28.90% | 13.93% | 18.18% | 7.98% | 2.48% | 5.01% | 3.24% | 1.45% |
| Zd Momo | 17.45% | 27.17% | 12.70% | 17.39% | 10.46% | 2.81% | 6.51% | 4.01% | 1.51% |
| Zd Gigi_4to1 | 19.84% | 30.22% | 14.63% | 19.16% | 6.57% | 1.89% | 3.89% | 2.55% | 1.24% |
| Zd Momo_4to1 | 19.19% | 29.72% | 14.06% | 19.08% | 7.63% | 1.80% | 4.45% | 2.79% | 1.28% |
| Td FL | 27.81% | 29.89% | 13.59% | 20.11% | 3.45% | 1.01% | 2.20% | 1.15% | 0.79% |
| Td KS | 24.16% | 26.85% | 12.12% | 18.04% | 10.14% | 0.98% | 5.56% | 1.27% | 0.88% |

**Supp. Table 3.**
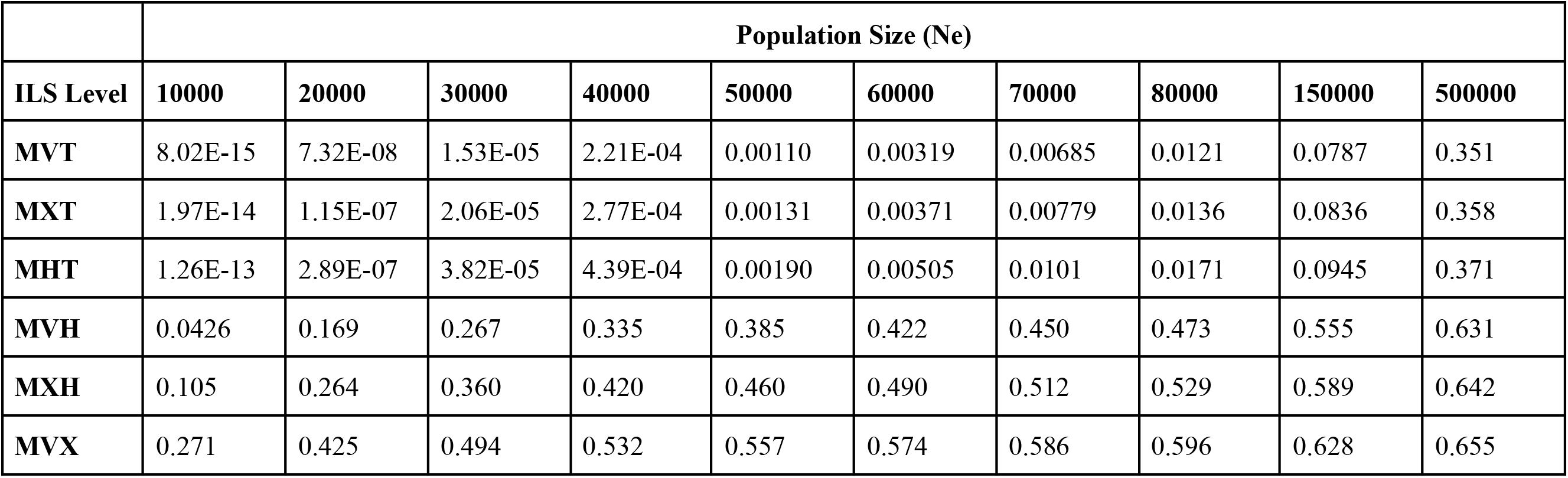
Probability of discordance due to incomplete lineage sorting across a range of divergence times (ILS Levels) and population sizes (Ne).

| | Population Size ($N_e$) | | | | | | | | | |
| --- | --- | --- | --- | --- | --- | --- | --- | --- | --- | --- |
| ILS Level | 10000 | 20000 | 30000 | 40000 | 50000 | 60000 | 70000 | 80000 | 150000 | 500000 |
| MVT | 8.02E-15 | 7.32E-08 | 1.53E-05 | 2.21E-04 | 0.00110 | 0.00319 | 0.00685 | 0.0121 | 0.0787 | 0.351 |
| MXT | 1.97E-14 | 1.15E-07 | 2.06E-05 | 2.77E-04 | 0.00131 | 0.00371 | 0.00779 | 0.0136 | 0.0836 | 0.358 |
| MHT | 1.26E-13 | 2.89E-07 | 3.82E-05 | 4.39E-04 | 0.00190 | 0.00505 | 0.0101 | 0.0171 | 0.0945 | 0.371 |
| MVH | 0.0426 | 0.169 | 0.267 | 0.335 | 0.385 | 0.422 | 0.450 | 0.473 | 0.555 | 0.631 |
| MXH | 0.105 | 0.264 | 0.360 | 0.420 | 0.460 | 0.490 | 0.512 | 0.529 | 0.589 | 0.642 |
| MVX | 0.271 | 0.425 | 0.494 | 0.532 | 0.557 | 0.574 | 0.586 | 0.596 | 0.628 | 0.655 |

## Supplemental Figures

**Supp. Figure 1:**
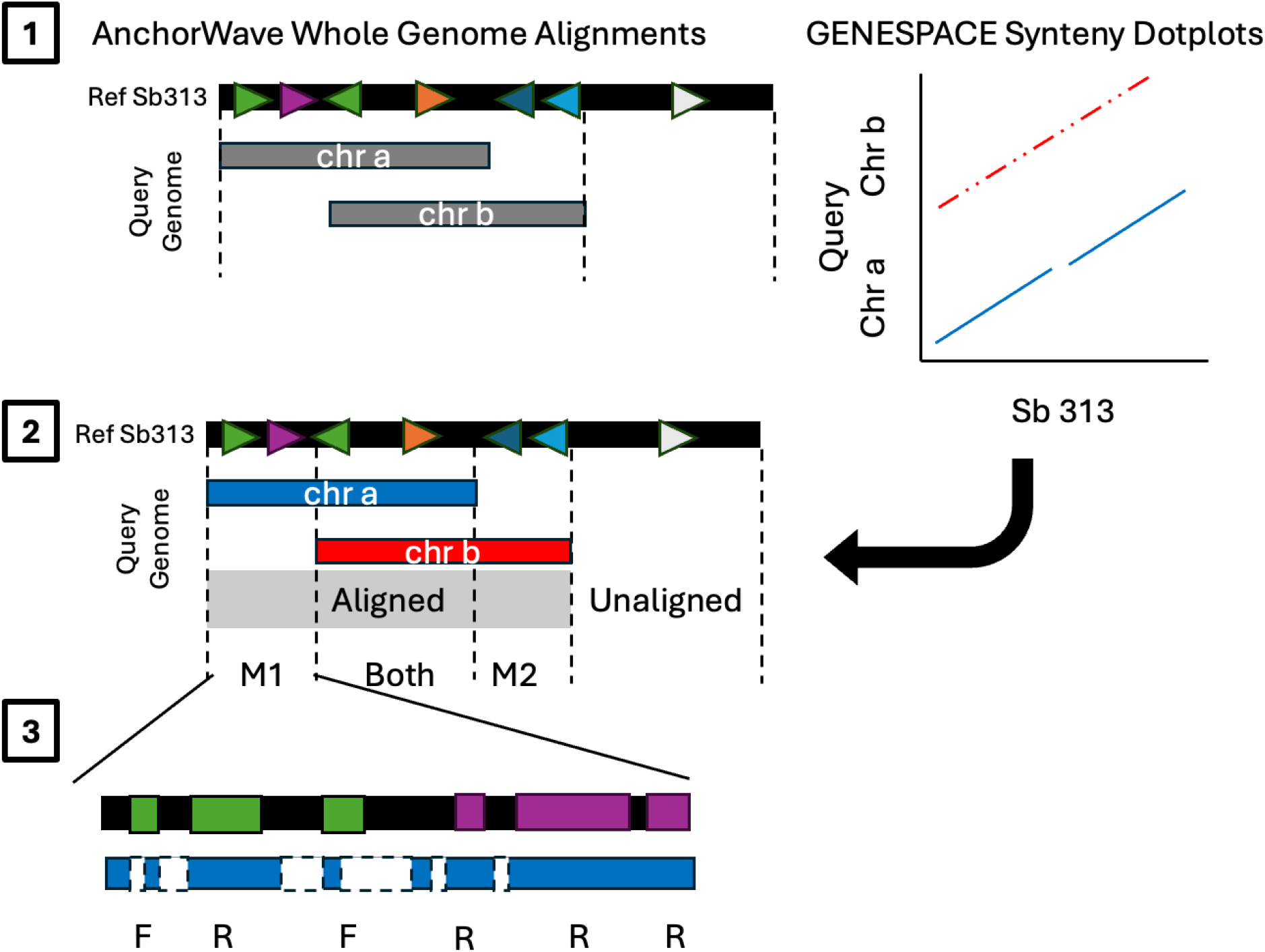
Diagram of how fractionation classifications are made using AnchorWave whole genome alignments and GENESPACE synteny dotplots. (1) Black rectangle represents the reference sorghum genome with different reference genes represented with colored triangles. Query Tripsacinae genome represented by grey rectangles where two chromosomes (a and b) have aligned to the same region in sorghum, though there are some areas where there’s no overlap. Dashed lines represent the boundaries where there is alignment between the query genome and the reference genome. From the synteny dotplot, we assign chr b to the M2 subgenome and chr a to the M1 subgenome. (2) The information is combined to segregate reference exons into those that are “aligned” vs “unaligned” by subgenome. (3) For each aligned sorghum reference exon (colored boxes), if a deletion (white box with dashed borders), is called in the query genome that overlaps the exon boundaries, the exon is called fractionated (F). If the exon is aligned and does not overlap with any deletion calls, it is called retained (R) .

**Supp. Figure 2.**
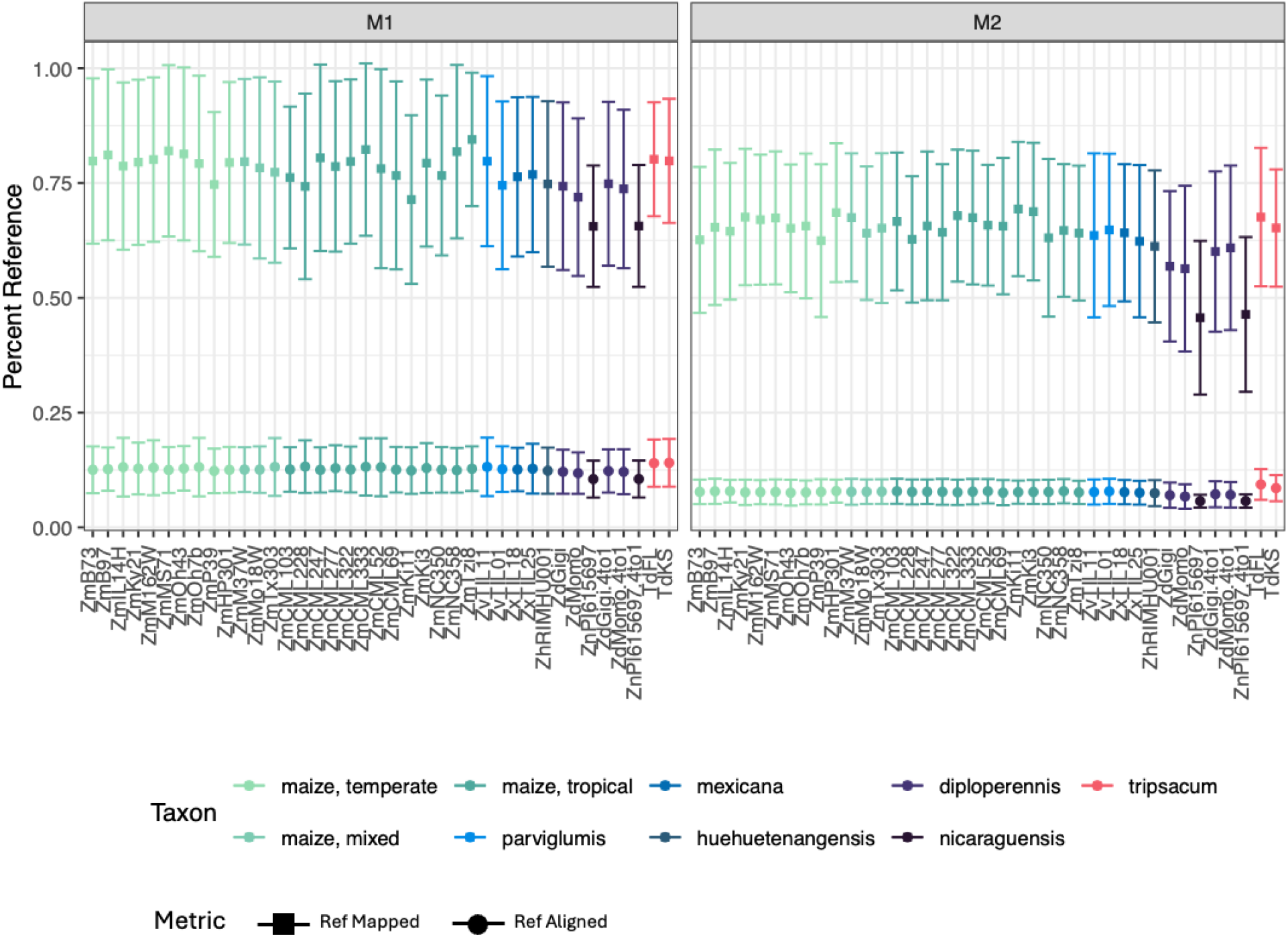
Output from VCFMetrics plugin on the genome-wide rate of mapping vs. alignment across genomes relative to sorghum, averaged across the 10 sorghum reference chromosomes, and split by subgenome (M1 and M2). Mapped (square points) refers to any matches (nucleotide to nucleotide or gap) while aligned (circle points) refers to exact matches (nucleotide to nucleotide). Thus, mapping rates are higher than alignment rates because of large amounts of deletion in Tripsacinae genomes relative to sorghum. Points correspond to the mean rate (1 = 100%) per genome and error bars represent 1 standard deviation. Points are colored according to broad taxonomic groups. *Diploperennis* and *nicaraguensis* genomes with “4to1” suffix indicate alignments which were run with 4:1 alignment parameters to account for unresolved haplotypes within the assemblies whereas those without the suffix use the standard 2:1 alignment parameters. Genome names refer to *Tripsacum dactyloides* (Td FL, Td KS), *Zea nicaraguensis* (Zn PI615697), *Zea diploperennis* (Zd Momo, Zd Gigi), *Zea mays* ssp. *huehuetenangensis* (Zh RIMHU001), *Zea mays* ssp. *mexicana* (Zx TIL25, Zx TIL18), *Zea mays* ssp. *parviglumis* (Zv TIL01, Zv TIL11), and *Zea mays* ssp. *mays* (Tropical: Zm Tzi8, Zm NC358, ZmNC350, Zm Ki3, Zm Ki11, Zm CML69, Zm CML52, Zm CML333, Zm CML322, Zm CML277, Zm CML247, Zm CML228, Zm CML103. Mixed: Zm Tx303, Zm Mo18W, Zm M37W. Temperate: Zm P39, Zm HP301, Zm Oh7b, Zm Oh43, Zm MS71, Zm M162W, Zm Ky21, Zm IL14H, Zm B97, Zm B73).

**Supp. Figure 3.**
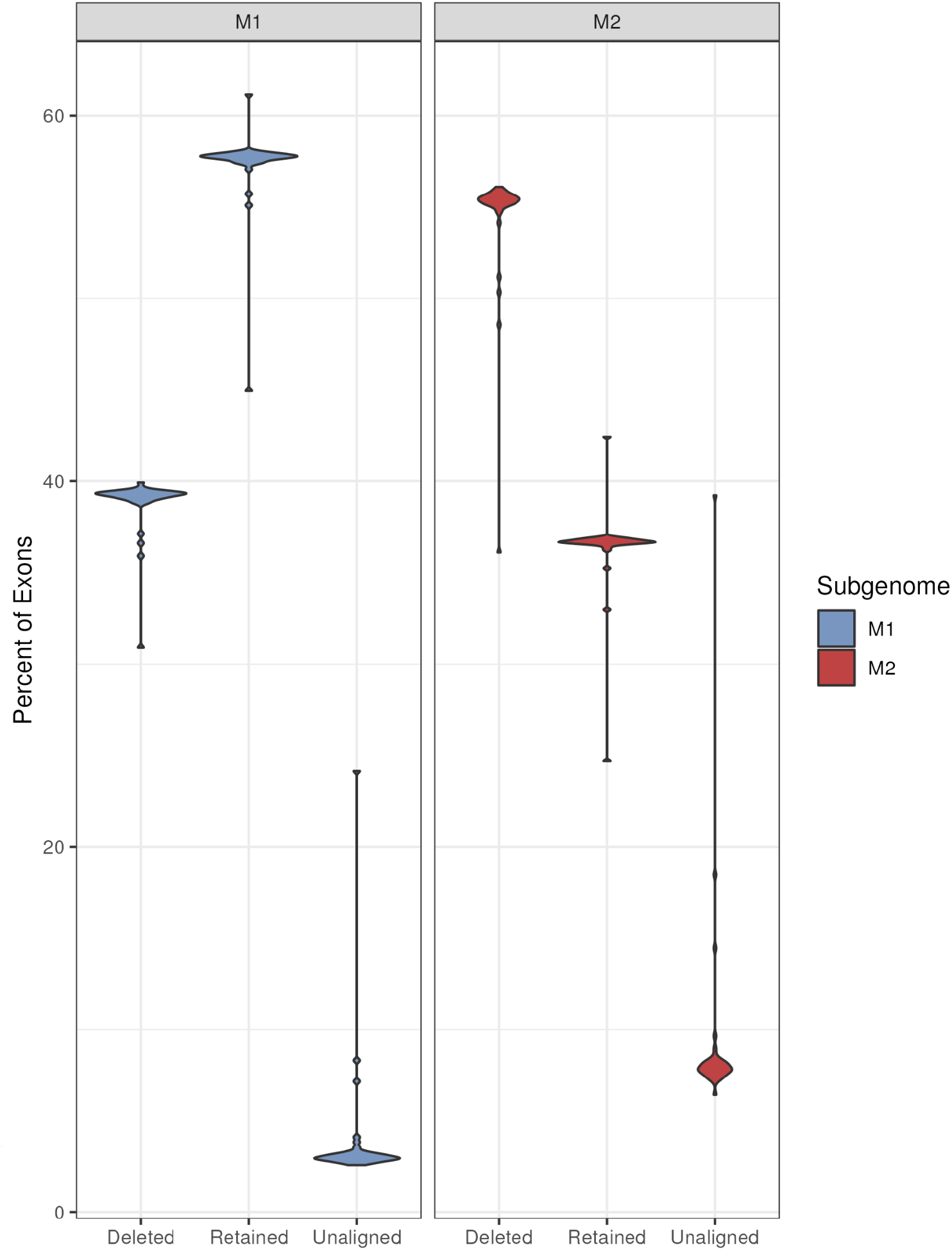
Percentage of reference exons identified as aligned and either completely retained (“Retained”) or some loss (“Deleted”) or as “Unaligned” across subgenomes M1 and M2 for all Tripsacinae genomes. These include the *Z. nicaraguensis* genome which corresponds to the long tail in the unaligned categories.

**Supp. Figure 4.**
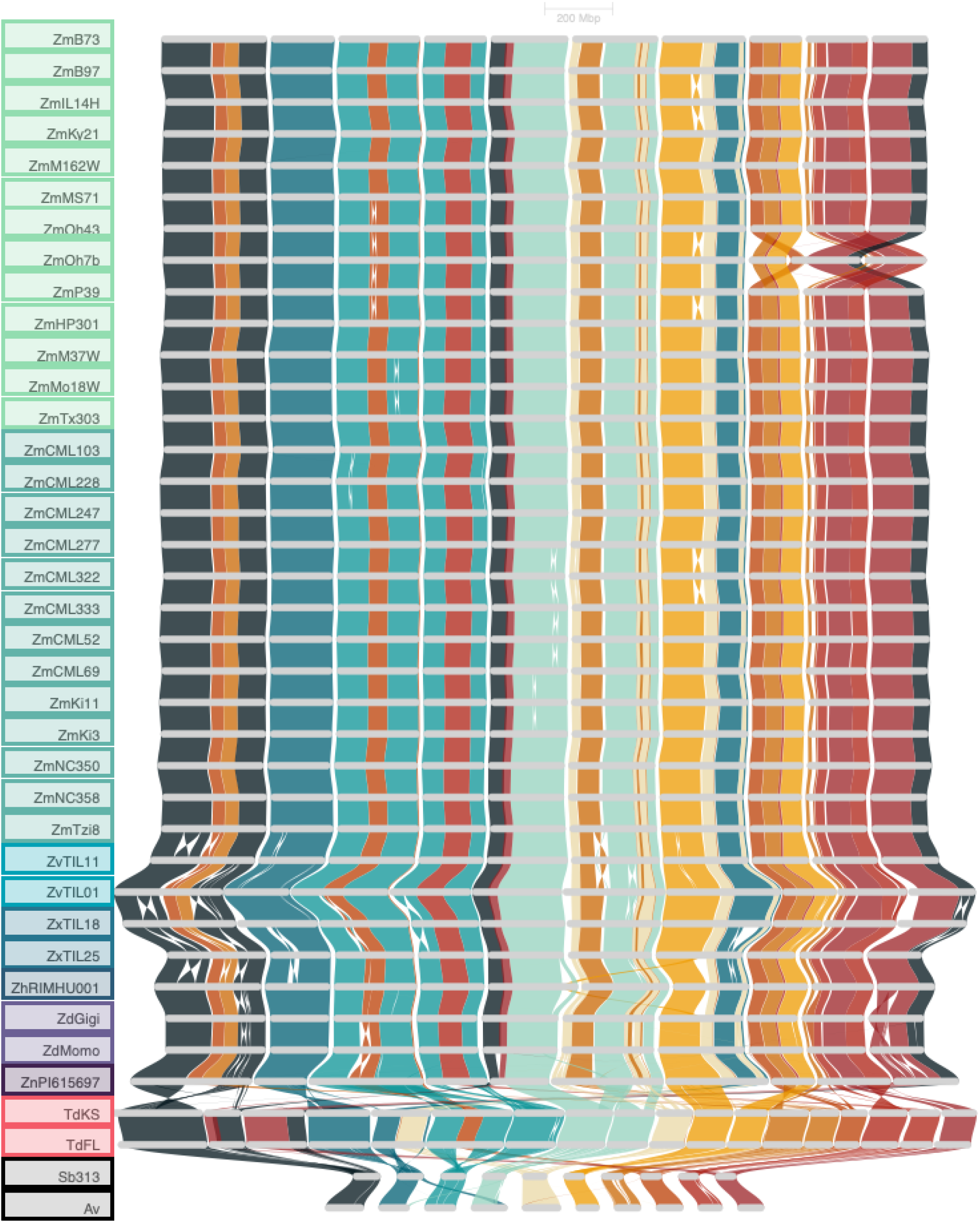
Riparian plot between all genomes. Colored braids show syntenic blocks between genomes, colored by sorghum reference chromosome going from chromosome 1 in dark blue to chromosome 10 in red. Twists in the colored braids indicate inversions between genomes. Genome species from the bottom to top: *Anatherum virginicum* (Av), *Sorghum bicolor* (Sb313), *Tripsacum dactyloides* (Td FL, Td KS), *Zea nicaraguensis* (Zn PI615697), *Zea diploperennis* (Zd Momo, Zd Gigi), *Zea mays* ssp. *huehuetenangensis* (Zh RIMHU001), *Zea mays* ssp. *mexicana* (Zx TIL25, Zx TIL18), *Zea mays* ssp. *parviglumis* (Zv TIL01, Zv TIL11), and *Zea mays* ssp. *mays* (Tropical: Zm Tzi8, Zm NC358, ZmNC350, Zm Ki3, Zm Ki11, Zm CML69, Zm CML52, Zm CML333, Zm CML322, Zm CML277, Zm CML247, Zm CML228, Zm CML103. Mixed: Zm Tx303, Zm Mo18W, Zm M37W. Temperate: Zm P39, Zm HP301, Zm Oh7b, Zm Oh43, Zm MS71, Zm M162W, Zm Ky21, Zm IL14H, Zm B97, Zm B73).

**Supp. Figure 5.**
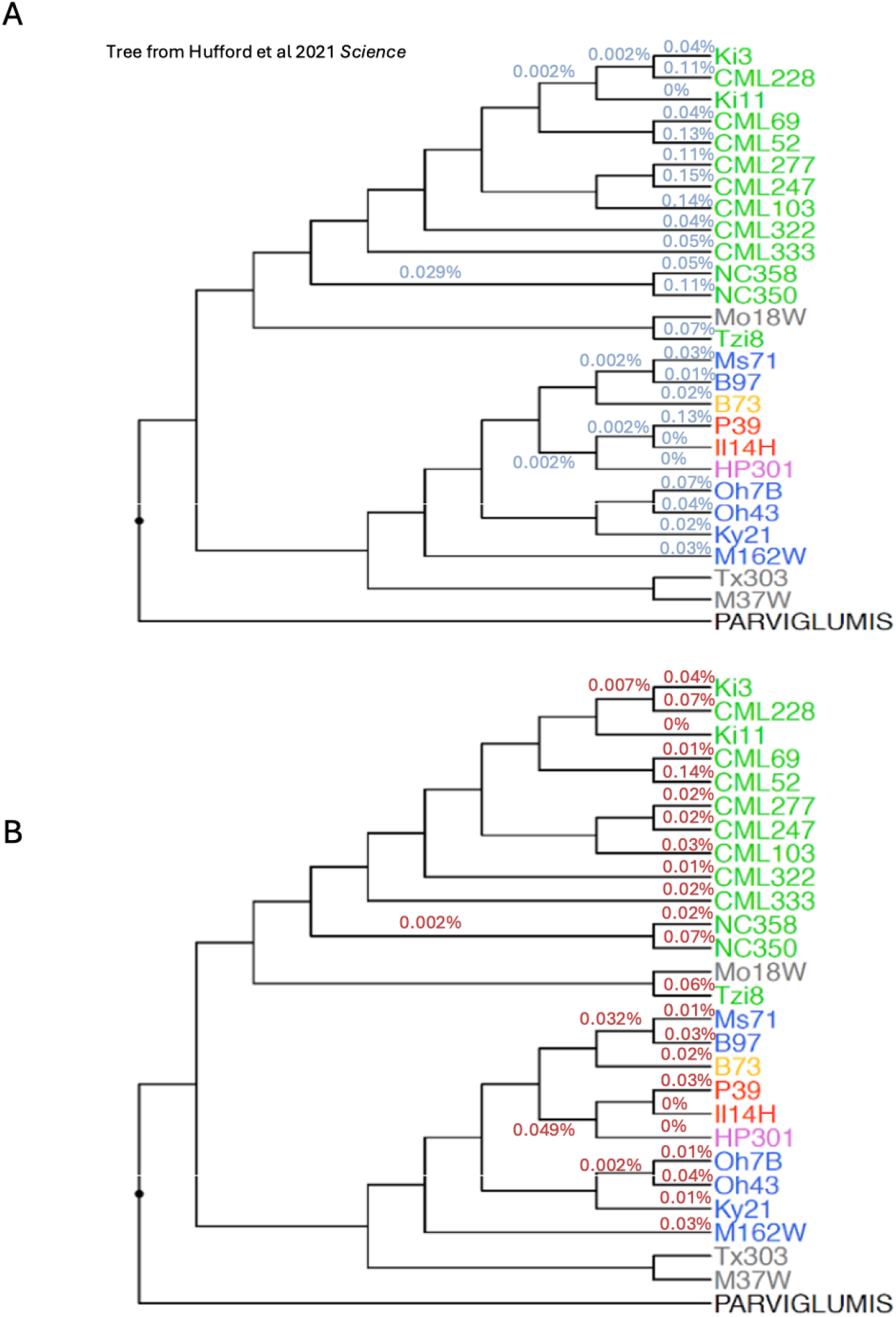
Sharing of fractionation across genomes with a focus on the maize NAM branches by subgenome, phylogeny from Hufford et. al. (2021). Exons must be aligned across all genomes to be included. Each branch is labeled with percent of exons in each subgnome that have been fractionated in every descendent lineage of the branch and retained in all other lineages. To reduce figure complexity, only internal branches with some specific fractionation are labeled. If a branch is unlabeled, there is no unique, monophyletic fractionation observed for that branch. Only NAM lines with tropical (green) or temperate only (non-stiff stalk = blue, stiff stalk = yellow, sweet corn = orange, and popcorn = pink) ancestry are used for estimates (mixed NAM lines (grey) which are products of recent admixture between temperate and tropical populations are ignored, Mo18W, Tx303, M37W).

**Supp. Figure 6.**
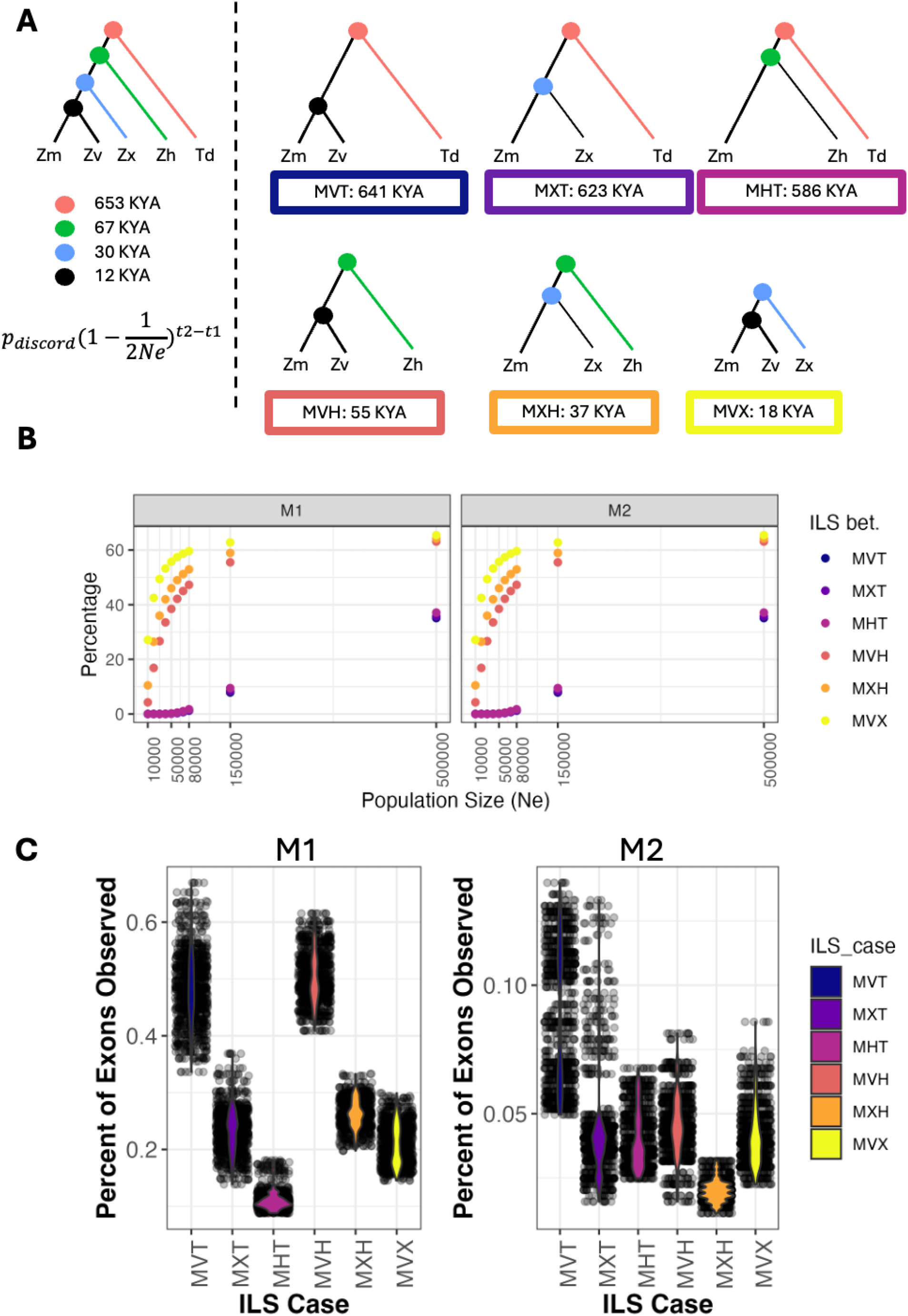
Expected vs. observed simple cases of discordance and ILS. **(A)** Cartoon diagrams of the various cases of discordance (ILS case) investigated. We use a simplified five taxon phylogeny (left) with the divergence times from Chen et al. (2022) as the backdrop for the rest of the analyses. We assume that 1 year equals 1 generation. Probability of expected discordance by ILS was calculated using the equation Pdiscord(1-1/2Ne)^t2-t1. Pdiscord refers to the proportion of trees that would be discordant. Since all cases we consider are a three lineage case, this is 2/3 for all cases. Each case is illustrated (right) with the label of the case and the t2-t1 value below the cartoon phylogeny. Outline colors of each case correspond across all panels. Cases are organized from the largest t2-t1 values to the smallest. **(B)** The expected percent of exons that should be discordant due to ILS. We multiply the probability calculated from (A) across a range of Ne (Chen et al 2022 plus 150K and 500K representing liberal overestimates of Ne) with the number of exons used for each subgenome to get the expected exon counts and then convert that to exon percentages. **(C)** The percentage of exons observed to be discordant according to each ILS case. One genome was picked randomly to represent each lineage and discordance was counted. This random sampling was repeated 1000 times to create a distribution. Note that the y-axis here is very small and the same units as (B).

**Supp. Figure 7.**
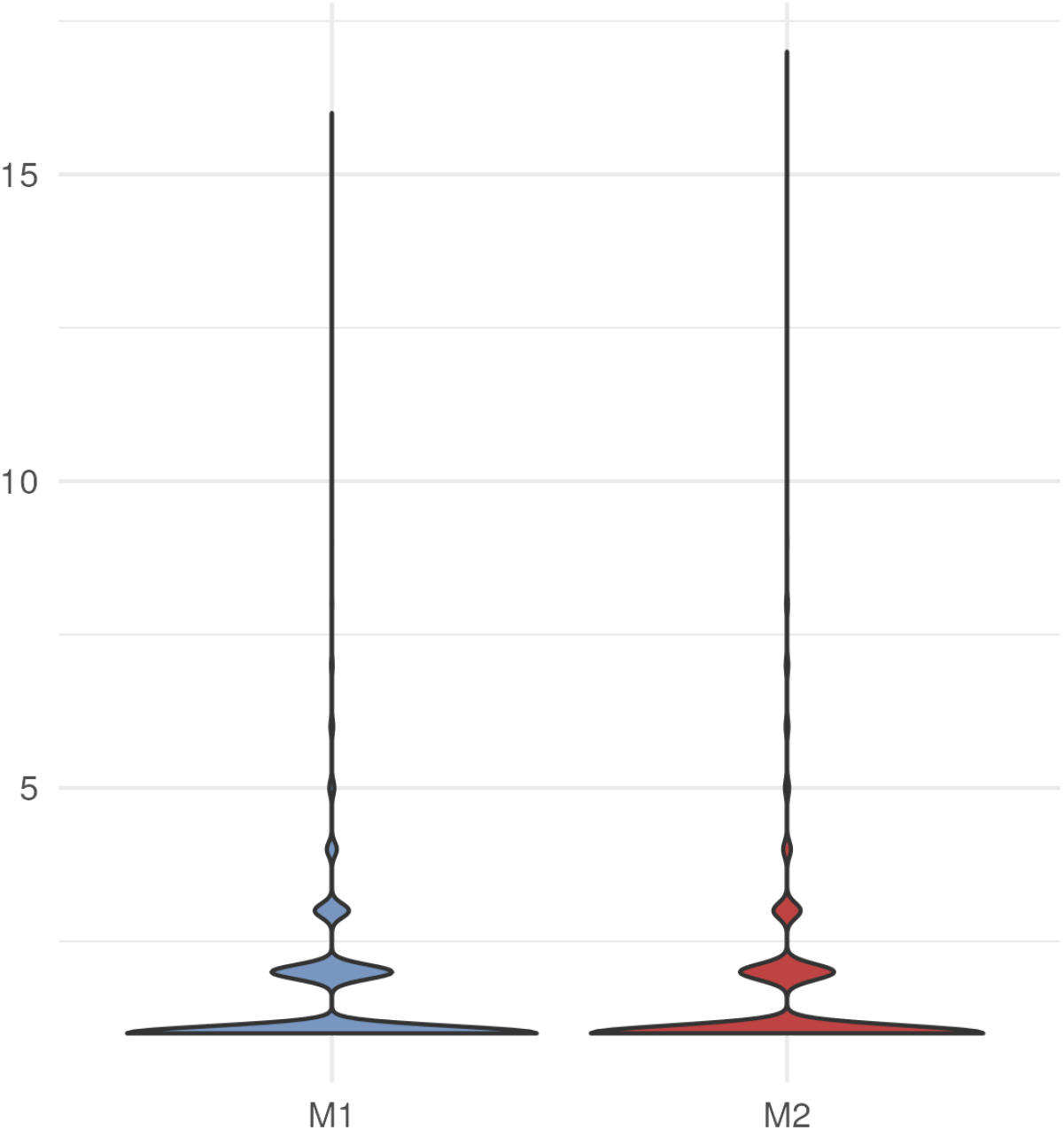
Number of ways a homoeolog was lost. Violin plot shows the distribution of homoeologs by subgenome that show different patterns of loss, starting at 1 loss pattern. Loss patterns are defined as the combination of exons called fractionated within the gene model.

**Supp. Figure 8.**
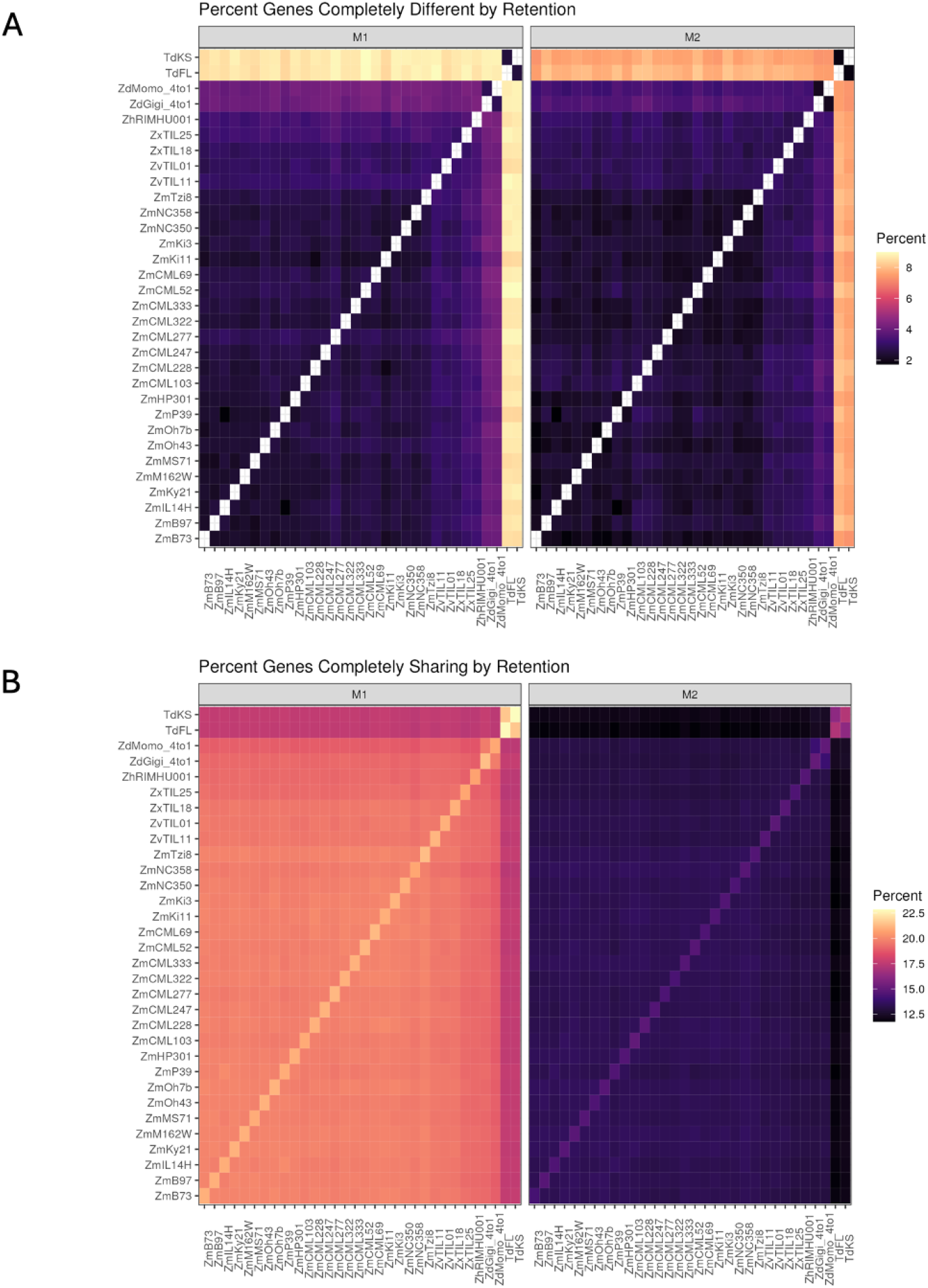
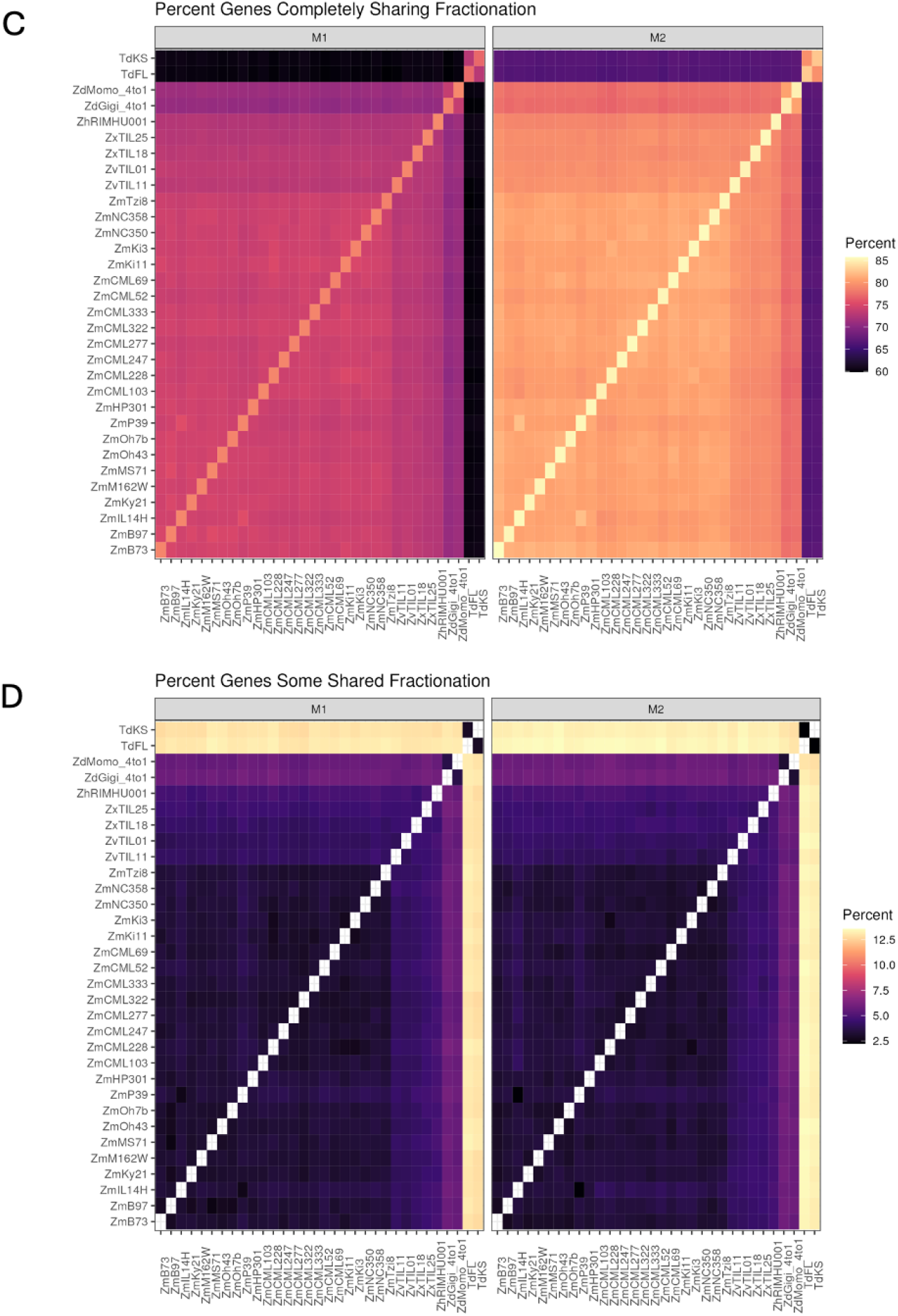
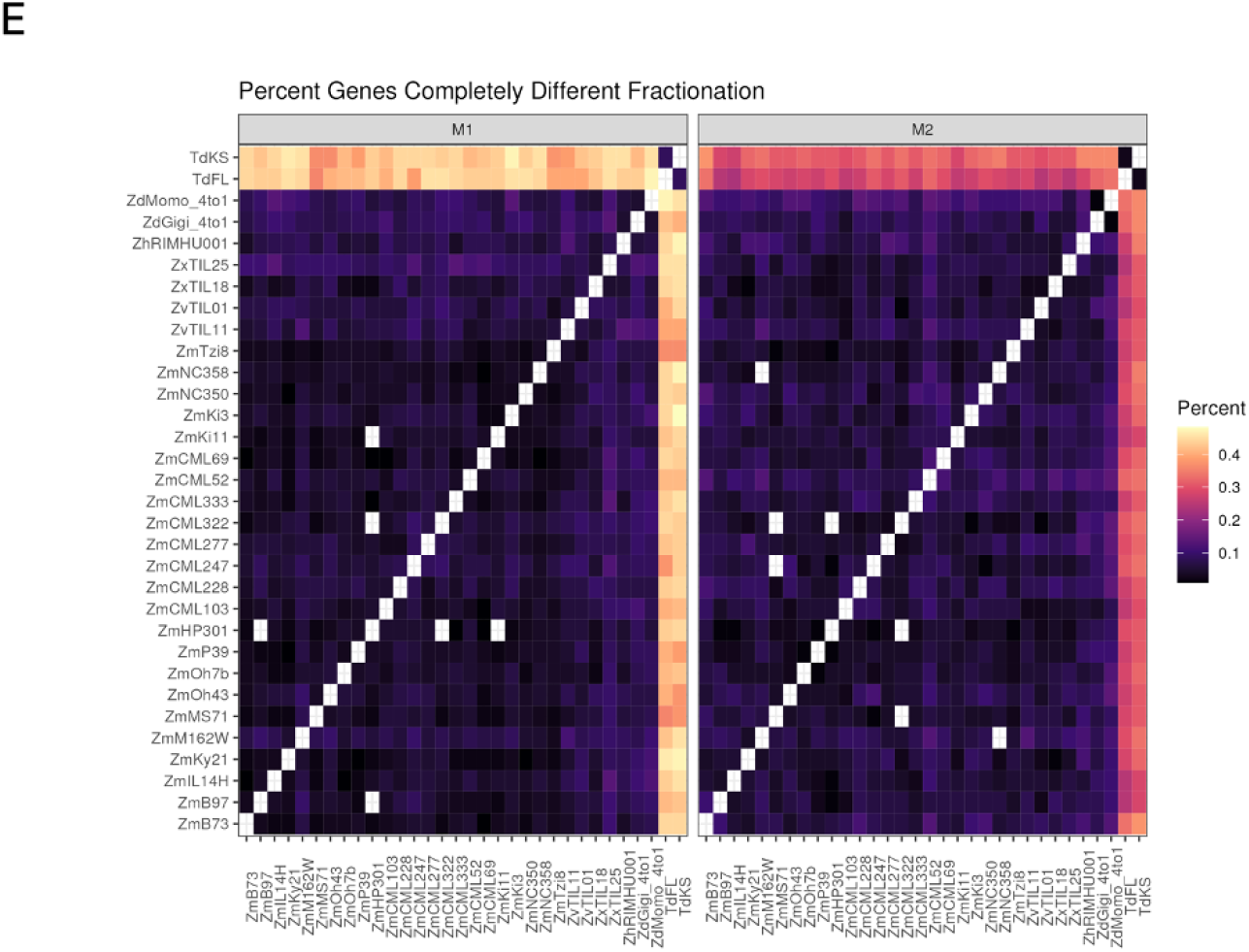
Heatmap showing the percentage of genes between any pair-wise comparison of genomes that show different levels of fractionation sharing. Empty tiles correspond to no observations of that category between those two genomes. Brighter colors indicate more genes. Genome names refer to *Tripsacum dactyloides* (Td FL, Td KS), *Zea diploperennis* (Zd Momo, Zd Gigi), *Zea mays* ssp. *huehuetenangensis* (Zh RIMHU001), *Zea mays* ssp. *mexicana* (Zx TIL25, Zx TIL18), *Zea mays* ssp. *parviglumis* (Zv TIL01, Zv TIL11), and *Zea mays* ssp. *mays* (Tropical: Zm Tzi8, Zm NC358, ZmNC350, Zm Ki3, Zm Ki11, Zm CML69, Zm CML52, Zm CML333, Zm CML322, Zm CML277, Zm CML247, Zm CML228, Zm CML103. Mixed: Zm Tx303, Zm Mo18W, Zm M37W. Temperate: Zm P39, Zm HP301, Zm Oh7b, Zm Oh43, Zm MS71, Zm M162W, Zm Ky21, Zm IL14H, Zm B97, Zm B73).

**Supp. Figure 9.**
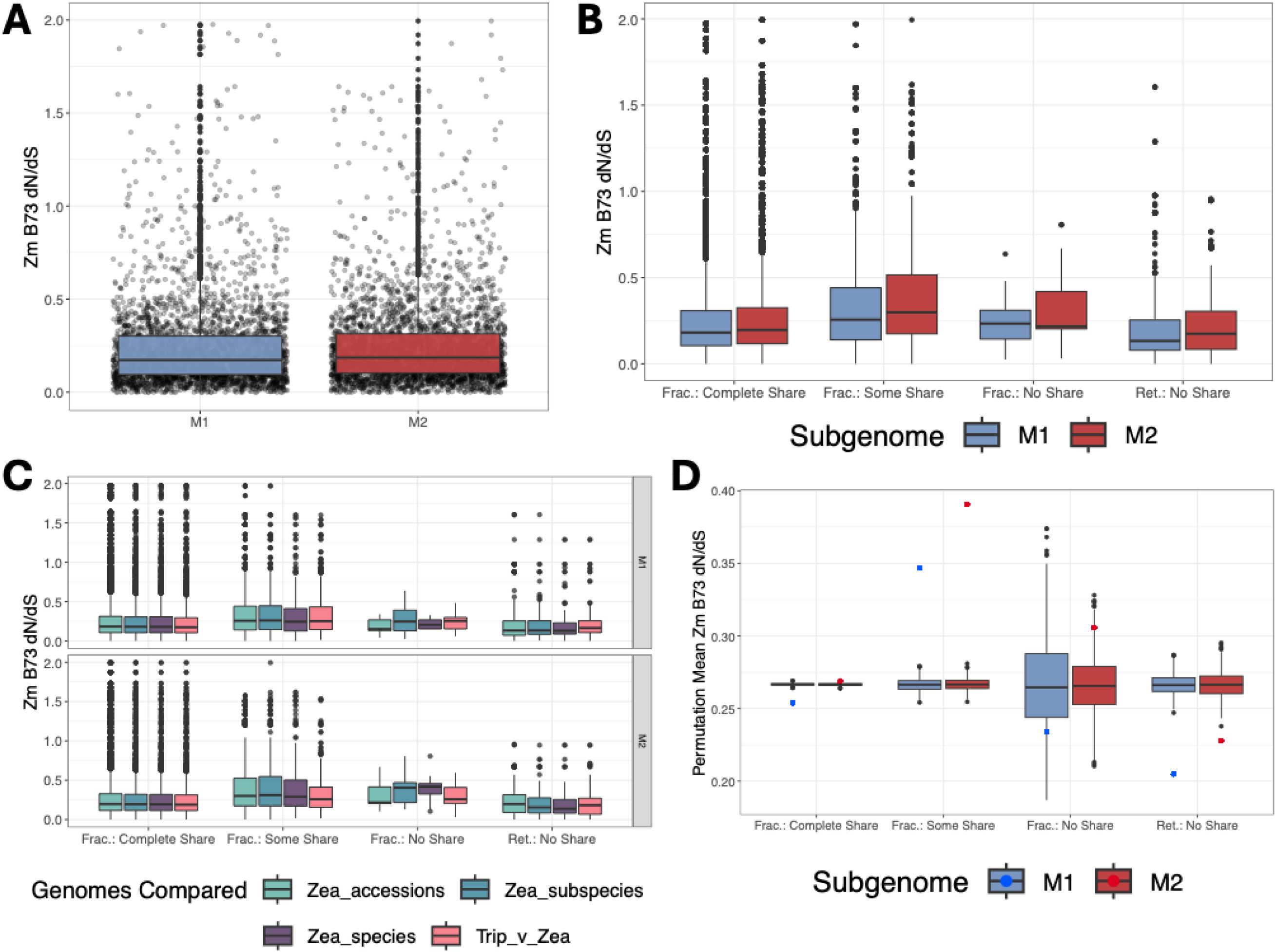
dN/dS from Yin et al. (2022) compared to fractionation sharing relative to Zm B73. (A) Overall dN/dS of the 3195 genes from Yin et al (2022) for Zm B73 split by subgenome. Boxplots colored by subgenomes with black points representing the individual dN/dS of each homoeolog. (B) Zm B73 dN/dS by sharing of the fractionation pattern between Zm B73 and all other genomes for each homoeolog. Boxplots colored by subgenomes. (C) Zm B73 dN/dS further subdivided by broad taxonomic differences between Zm B73 and the other genomes in addition to subgenome and fractionation pattern sharing. Boxplots are colored by taxonomic differences between Zm B73 and the other genomes. (D) Results of a permutation test where the boxplots represent the distribution of average dN/dS for 1000 permutations and colored points represent the observed average dN/dS for each category. Colors are by subgenome.

**Supp. Figure 10.**
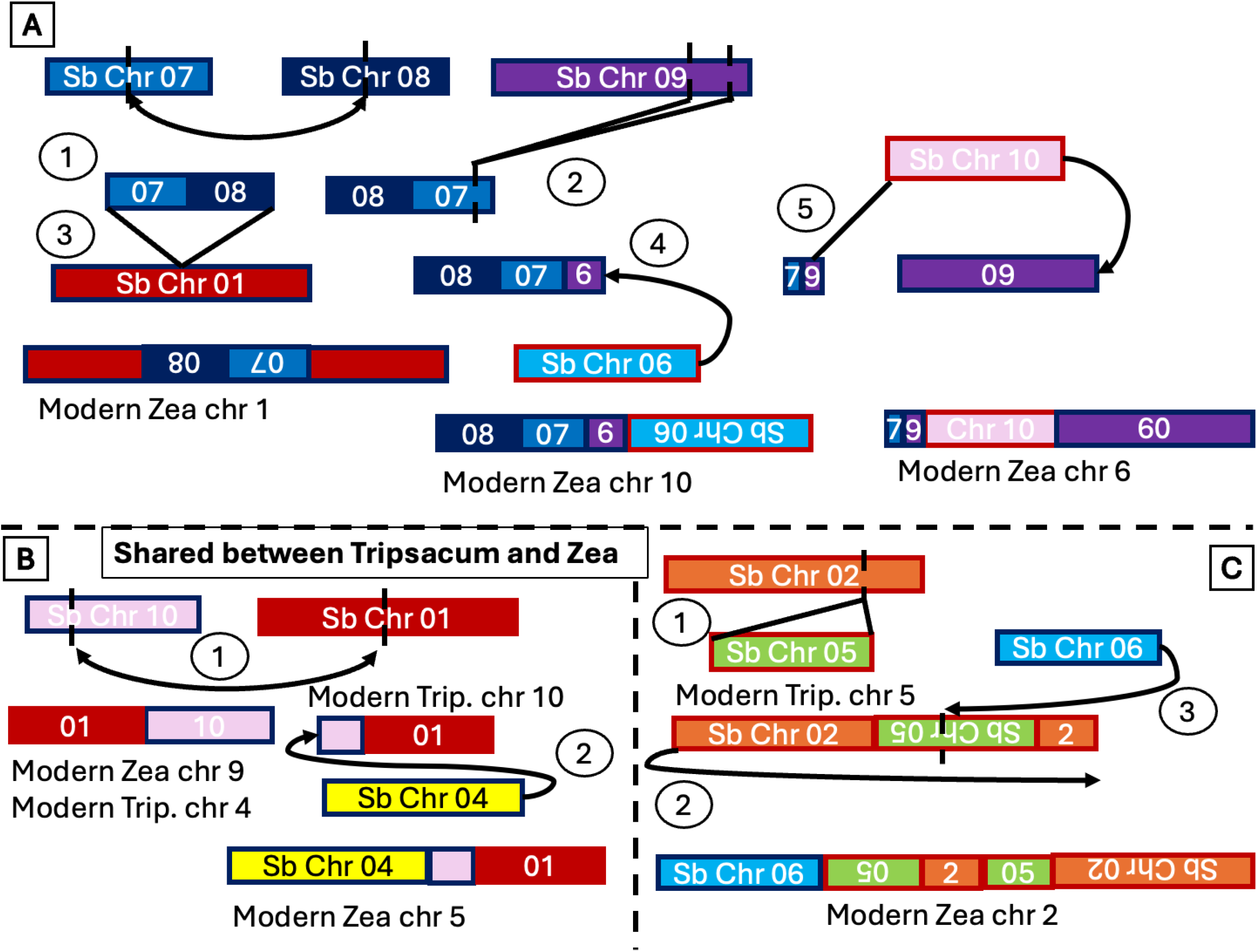
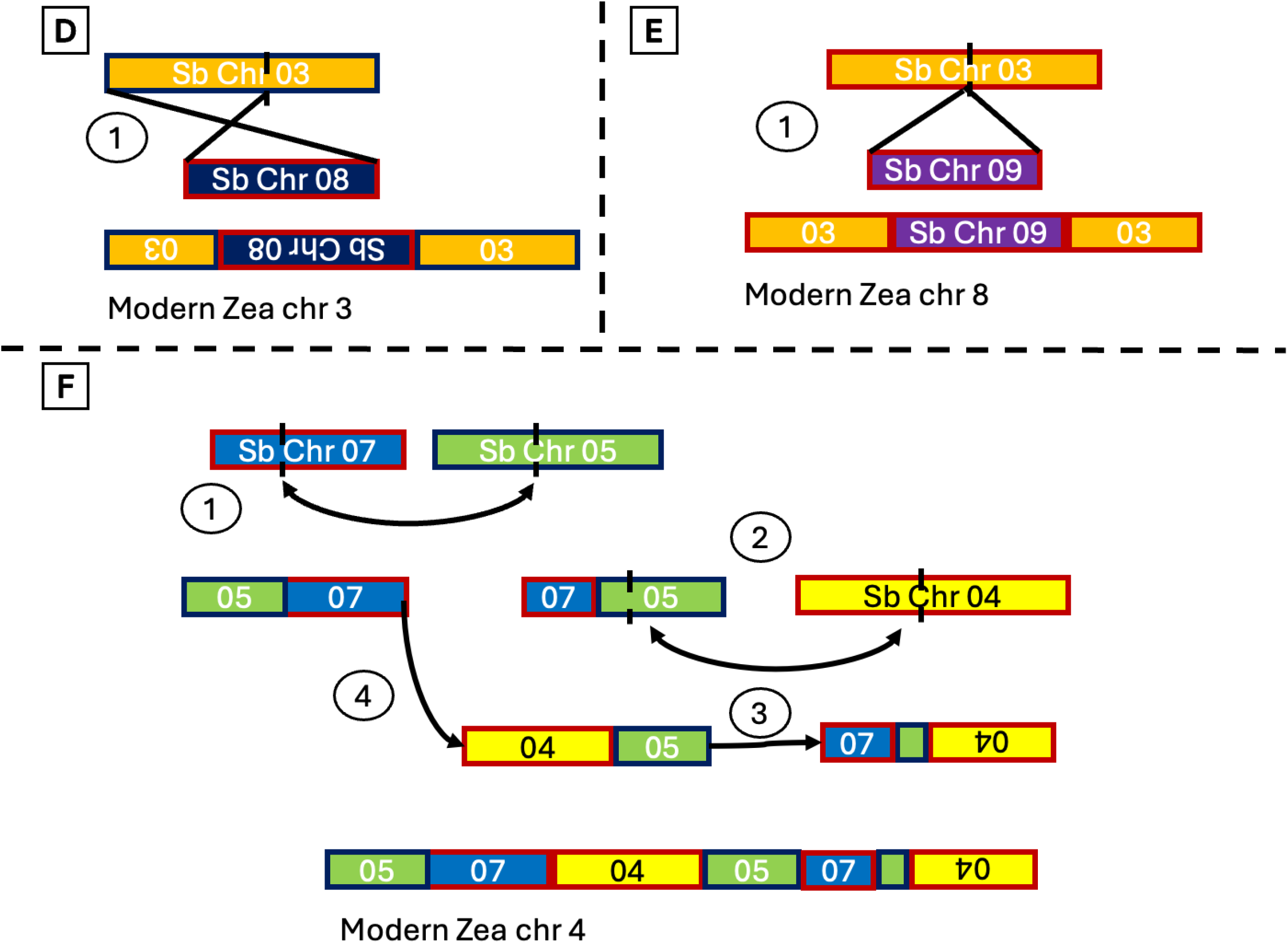
The modern *Zea* chromosomes may be the product of 6 series of reciprocal translocations and chromosomal fusions between ancestral chromosomes. Two of which are in part shared with *Tripsacum*. These figures represent simplified hypotheses based on syntenic dot plots of how these ancestral chromosomes rearranged to become the chromosomes observed today. The box fill color corresponds to modern sorghum chromosomes. The box outline colors correspond to ancestral subgenomes (blue = M1 and red = M2). **Series A)** (1) Reciprocal translocation between M1 chr 8 and M1 chr 7. (2) Reciprocal translocation between 8-7 segment and M1 chr 9 which also fissions a larger section of M1 chr 9. (3) The 7-8 segment inserts itself into M1 chr 1 resulting in the modern *Zea* chromosome 1. (4) The 8-7-9 segment fuses to M2 chr 6 resulting in the modern *Zea* chromosome 10. (5) The 7-9 segment and larger remaining segment of chr 9 fuse to M2 chr 10, resulting in the modern *Zea* chromosome 6. **Series B)** (1) Reciprocal translocation between M1 chr 10 and M2 chr 1. The 1-10 segment results in modern *Zea* chr 9 and modern *Tripsacum* chr 4. The 10-1 segment results in modern *Tripsacum* chromosome 10. (2) M1 chr 4 fuses to the end of the 10-1 segment resulting in modern *Zea* chromosome 5. **Series C)** (1) M2 chr 5 inserts into M2 chr 2, resulting in modern *Tripsacum* chromosome 5. (2) The 2-5-2 proto-chromosome fissions and recombines. (3) M1 chr 6 fuses to the end of the 5-2-5-2 proto-chromosome resulting in modern *Zea* chromosome 2. **Series D)** (1) M2 chromosome 8 inserts into M1 chromosome 3, resulting in modern *Zea* chromosome 3. **Series E)** (1) M2 chromosome 9 inserts into M2 chromosome 3, resulting in modern *Zea* chromosome 8. **Series F)** (1) Reciprocal translocation between M2 chr 7 and M1 chr 5. (2) Reciprocal translocation between the 7-5 segment and M2 chr 4. (3) Fusion of the segments from (2). (4) Fusion of the 5-7 segment from (1) to the 4-5-7-5-4 segment from (3), resulting in the modern *Zea* chromosome 4.

